# Parameter quantification for oxygen transport in the human brain

**DOI:** 10.1101/2024.04.13.589308

**Authors:** Yun Bing, Tamás I. Józsa, Stephen J. Payne

**Affiliations:** Institute of Biomedical Engineering, Department of Engineering Science, University of Oxford, Oxford, UK; Centre for Computational Engineering Sciences, School of Aerospace, Transport and Manufacturing, Cranfield University, Cranfield, UK; Institute of Applied Mechanics, National Taiwan University, Taipei, Taiwan

## Abstract

Oxygen is carried to the brain by blood flow through generations of vessels across a wide range of length scales. This multi-scale nature of blood flow and oxygen transport poses challenges on investigating the mechanisms underlying both healthy and pathological states through imaging techniques alone. Recently, multi-scale models describing whole brain perfusion and oxygen transport have been developed. Such models rely on effective parameters that represent the microscopic properties. While parameters of the perfusion models have been characterised, those for oxygen transport are still lacking. In this study, we set to quantify the parameters associated with oxygen transport and their uncertainties. We first present a multi-scale, multi-compartment oxygen transport model based on a porous continuum approach. We then determine the effective values of the model parameters. By using statistically accurate capillary networks, geometric parameters (vessel volume fraction and surface area to volume ratio) that capture the microvascular topologies are found to be 1.42% and 627 [mm^2^/mm^3^], respectively. These values compare well with those obtained from human and monkey vascular samples. In addition, maximum consumption rates of oxygen are optimised to uniquely define the oxygen distribution over depth. Simulation results from a one-dimensional tissue column show qualitative agreement with experimental measurements of tissue oxygen partial pressure in rats. We highlight the importance of anatomical accuracy through simulation performed within a patient-specific brain mesh. Finally, one-at-a-time sensitivity analysis reveals that the oxygen model is not sensitive to most of its parameters; however, perturbations in oxygen solubilities and plasma to whole blood oxygen concentration ratio have a considerable impact on the tissue oxygenation. These findings demonstrate the validity of using a porous continuum approach to model organ-scale oxygen transport and draw attention to the significance of anatomy and certain parameter values.

## 1 Introduction

Oxygen, an essential substance for maintaining the normal functioning of aerobic cells, is transported to the brain via blood flow through generations of vessels whose diameters cover a wide range of length scales. A constant supply of oxygen to the brain is particularly important due to the tight coupling between tissue perfusion and metabolism. Such coupling is regulated by both global and local mechanisms including cerebral autoregulation and neurovascular coupling [1]. Any disruption in oxygen delivery that persists for more than just a few minutes can result in irreversible damage to the brain parenchyma. It is, therefore, important to understand the interplay between these mechanisms that occur at different length scale.

Imaging techniques, such as PET (positron emission tomography) and BOLD-fMRI (blood-oxygenation level dependent functional magnetic resonance imaging), have provided invaluable insights on changes in cerebral haemodynamics and oxygenation in both healthy [2–4] and pathological [5–7] states. However, quantitative data produced by these techniques are often an indirect measure of cerebral oxygenation and metabolism, especially for the values of oxygen partial pressure 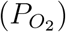. Although the cerebral microcirculation is the primary site for oxygen exchange and thus plays a crucial role in meeting the energy requirements of the brain, the spatial resolution (in the order of millimetres) of these techniques for *in vivo* imaging of the human brain is not high enough to reveal the influence of the microvasculature on perfusion and oxygenation.

These shortcomings have motivated the development of mechanistic models, notably at the micro-scale, to analyse as well as quantify perfusion and oxygen transport. Such micro-scale models employ discrete vascular networks either reconstructed directly from cerebral angiography [8–11] or computationally generated based on microvascular statistics [12–15]. Although these network models are useful tools in understanding localised interactions between blood flow patterns and oxygen distribution, not only are they difficult to validate but it is computationally intensive to scale up to the organ-scale. On the other hand, whole brain simulations are mostly based on lumped compartments, such as the works of [16–18]. These models are highly simplified without taking into account the cerebral anatomy or spatial variations within the compartments.

Multi-scale modelling is a promising method for describing complex biological systems that span across different length scales. Such models are able to represent the entire system while maintaining a reasonable computational efficiency. Homogenisation techniques are often used to construct multi-scale models, where governing equations at the macroscale are characterised by parameters computed based on the microscopic properties. Building upon the work of El-Bouri and Payne [12, 13], Józsa *et al*. [19–21] developed a three-dimensional (3D) perfusion model at the brain scale. This model is connected to a 1D large arterial model that describes blood flow from the heart to the leptomeningeal circulation [22, 23], enabling the study of pathological conditions such as ischaemic stroke. Recently, Payne and Mai [24] extended the perfusion model to account for changes in blood (vessel) volume and to include oxygen transport.

An important part of multi-scale modelling is to find effective parameter values that represent the microscopic properties. This is not a trivial task, especially for human physiological models, due to the lack of data and the gap between clinical measurements and mathematical representations. For perfusion models, the micro-scale heterogeneities are captured by the permeability tensors which have been characterised by El-Bouri and Payne [12] and Józsa *et al*. [19, 21]. For oxygen transport, geometric parameters (vessel volume fraction and surface area to volume ratio) encapsulate the microvascular topologies. Payne and Mai only estimated these parameter values based on orders of magnitude [24]. Another important yet often neglected aspect is sensitivity analysis. Through quantifying parameter uncertainties, key pathways within the model can be identified so are the potential simplifications that can be made on the model.

In this paper, we present a continuum-based porous multi-scale, multi-compartment oxygen model that is coupled to the aforementioned perfusion model developed by Józsa *et al*. [19, 20]. We then systematically estimate the parameters of the oxygen transport model. In particular, effective values of the geometric parameters are obtained using statistically accurate microvascular networks; and the maximum consumption rate of oxygen is optimised to uniquely define the oxygen distribution with respect to the depth of the brain. Next, we implement the model with a 1D tissue column and a 3D patient-specific brain mesh. We validate the 1D model against tissue 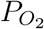 measured in rats and show the importance of anatomy accuracy using the 3D model. Finally, a sensitivity analysis is performed to quantify the uncertainties of the parameter values and identify the parameters to which the model results are most sensitive.

## 2 Model formulation

Figure 1 shows a schematic overview of the coupled model. In line with the perfusion model, the cerebral microvasculature below the cortical surface is divided into three discrete compartments according to their geometrical characteristics and their roles in oxygen transport. These three compartments are the penetrating arterioles, capillaries, and penetrating venules. Additionally, a fourth compartment concerning tissue (including the interstitial space) is included. The microvascular compartments receive oxygen from the macrocirculation through bulk flow of blood, i.e., advection (Process 1–3), and the arteriovenous diffusion shuts (Process 4) which refer to the phenomenon that oxygen leaving the arteries by diffusion enters the nearby veins. Oxygen is supplied to the tissue via diffusion across the vessel wall (Process 5–7), and it is then metabolised in the tissue compartment (Process 8).

**Figure 1.**
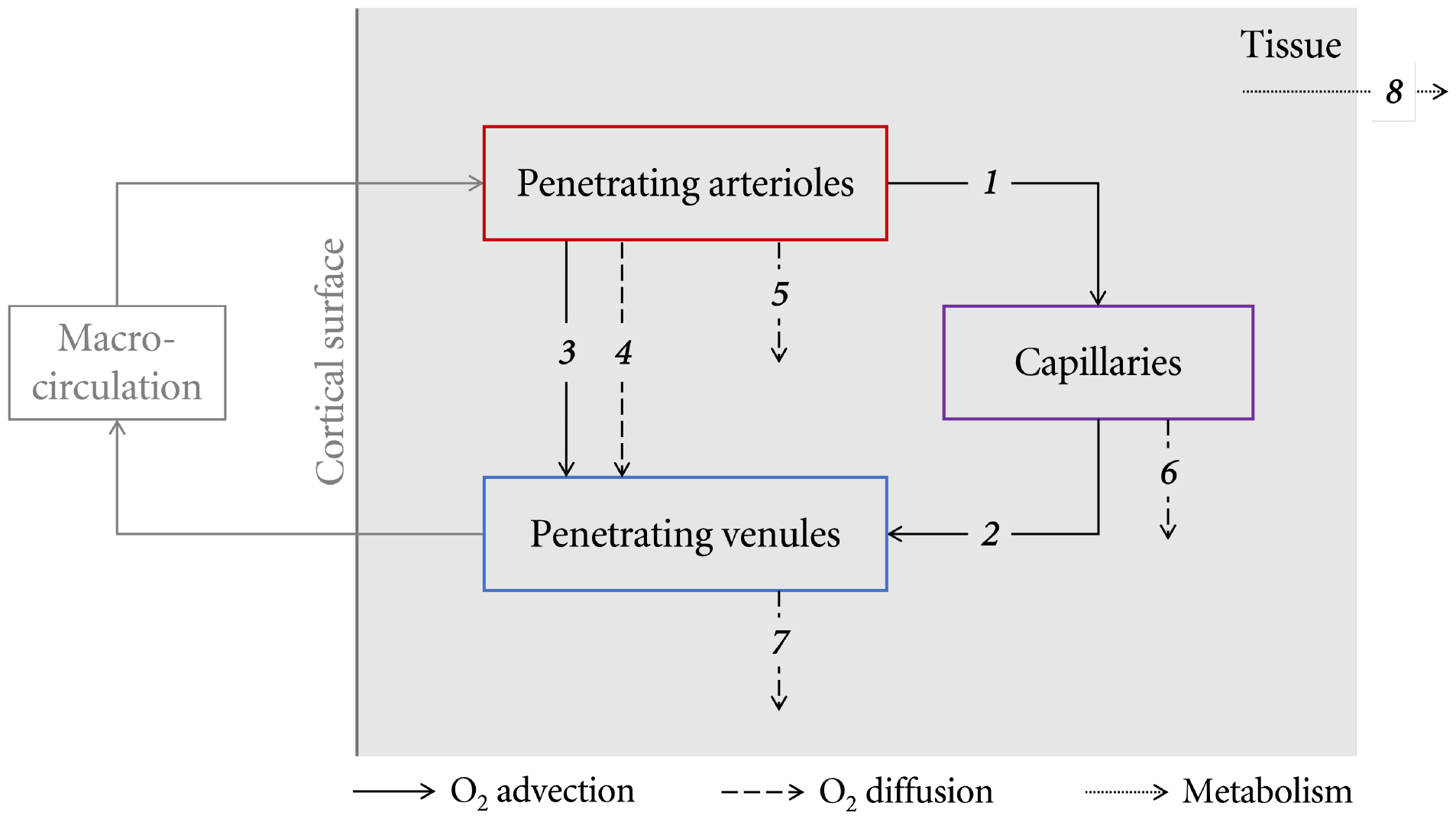
Illustration of inter-compartment oxygen transport processes. The arrows indicate the direction of net oxygen exchange.

Because of the complex nature of oxygen transport, a number of assumptions are made here to simplify the model. First of all, we assume that there is no direct oxygen transport, either through anastomoses (Process 3) or diffussion shuts (Process 4), from arterioles to venules. Even though the former’s presence is clearly observable under pathological conditions [25] and the latter has been observed in rats [26], there is a lack of evidence supporting the existence of such phenomenon in healthy humans [27–29]. Secondly, we eliminate Process 7 due to the small oxygen partial pressure difference between venules and tissue. This leaves the venular compartment to be essentially an oxygen sink. Lastly, we assume that blood is well-mixed and the radial concentration gradient within the vessel is negligible. The ‘non-well-mixed’ effect can be important when microscale blood flow is quantified [30]; however, the present study focuses on organ-scale modelling.

After applying the above assumptions, the oxygen transport model is derived directly on the macro-scale based on mass conservation; details of the derivation can be found in [31]. The resulting governing equations read as:

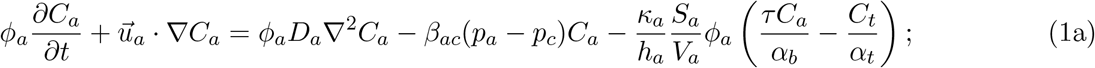

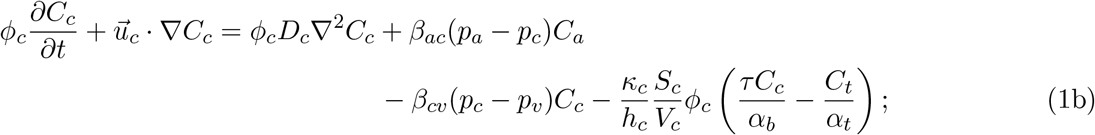

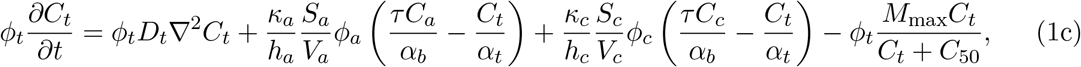

Here, *t* is time and subscript *a, c, v* and *t* represent arterioles, capillaries, venules and tissue, respectively. The unknown of the system is the oxygen concentration *C*_*a*_, *C*_*c*_ [mm^3^ O_2_/mm^3^ blood] and *C*_*t*_ [mm^3^ O_2_/mm^3^ tissue]. Parameters of the system can be categorised as either perfusion-related or oxygen-related; the meaning of each parameter and its unit are listed in Table 1.

**Table 1:**
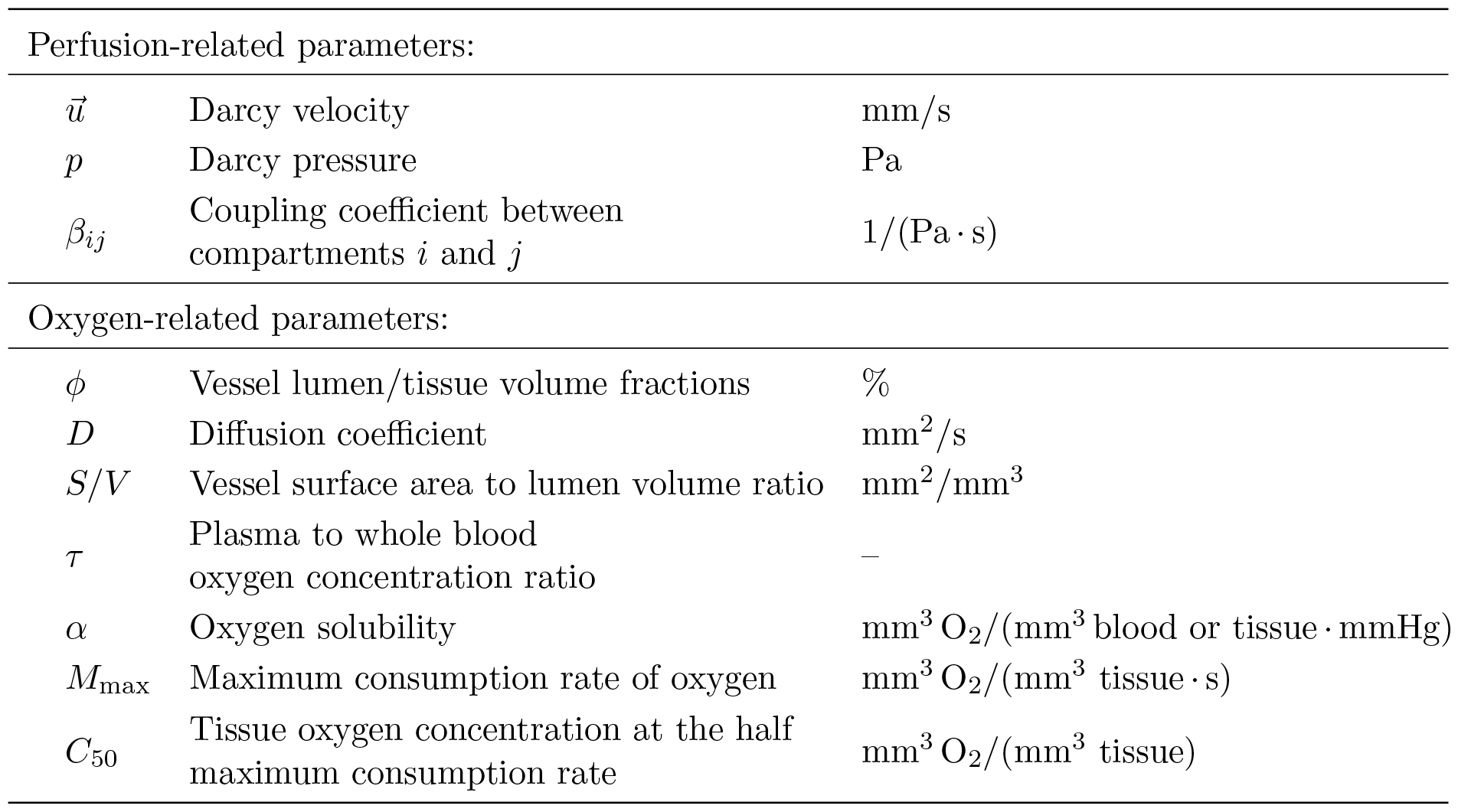
Meaning and units of the model parameters.

| Perfusion-related parameters: |  |  |
| --- | --- | --- |
| $\vec{u}$ | Darcy velocity | mm/s |
| $p$ | Darcy pressure | Pa |
| $\beta_{ij}$ | Coupling coefficient between compartments $i$ and $j$ | 1/(Pa · s) |
| Oxygen-related parameters: |  |  |
| $\phi$ | Vessel lumen/tissue volume fractions | % |
| $D$ | Diffusion coefficient | $\text{mm}^2/\text{s}$ |
| $S/V$ | Vessel surface area to lumen volume ratio | $\text{mm}^2/\text{mm}^3$ |
| $\tau$ | Plasma to whole blood oxygen concentration ratio | — |
| $\alpha$ | Oxygen solubility | $\text{mm}^3 \text{O}_2/(\text{mm}^3 \text{ blood or tissue} \cdot \text{mmHg})$ |
| $M_{\max}$ | Maximum consumption rate of oxygen | $\text{mm}^3 \text{O}_2/(\text{mm}^3 \text{ tissue} \cdot \text{s})$ |
| $C_{50}$ | Tissue oxygen concentration at the half maximum consumption rate | $\text{mm}^3 \text{O}_2/(\text{mm}^3 \text{ tissue})$ |

The physical significance of each term, taking Equation (1a) as an example, from left to right is: the local change of oxygen concentration; advection within the compartment; diffusion within the compartment; inter-compartment advection term; and inter-compartment diffusion term. Additionally, the last term in Equation (1c) is the metabolic term described by the widely-used Michaelis–Menten kinetics [32].

The boundary conditions imposed with Equation (1) in a computational domain Ω are

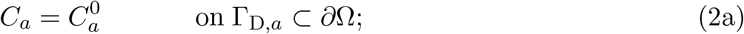

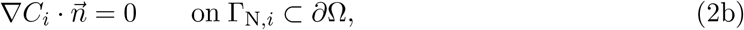

with *i* indicating *a, c* and *t*, ∂Ω representing the boundary surface of Ω and 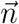 being the outward-pointing unit vector perpendicular to ∂Ω. The Dirichlet boundary condition (DBC), Equation (2a), indicates that a value of 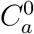 is prescribed to *C*_*a*_ on the cortical surface of the arteriole compartment (Γ_D,*a*_). The homogenous Neumann boundary condition (NBC), Equation (2b), ensures that no oxygen crosses the boundary Γ_N_ = ∂Ω \ Γ_D_.

However, one-dimensional analysis performed by Bing [31] showed that only less than half of the oxygen at any given depth was being delivered towards the deeper layer of the arteriolar compartment and that oxygen was predominantly transported into the venules before it was able to distribute within the capillary compartment, which led to insufficient oxygenation in the white matter. This is due to the relatively large inter-compartment advection effects and that the only source of oxygen is from the cortical surface.

Although large decrease in partial pressure of oxygen in arterioles 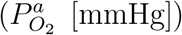 was observed with depth in anaesthetised animals, indicating a directly contribute to tissue oxygenation from arterioles [33, 34], small to no variation in 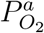 was detected along penetrating arterioles in awake mice [35, 36]. Morphologically speaking, there are arterioles that exclusively vascularise the white matter [29]. The inclusion of such arterioles as an additional compartment would allow the white matter to be better oxygenated through a more direct pathway. Unfortunately, the only statistical information available regarding these vessels is their diameter range, which is not enough for model construction. Instead, we assume that there is negligible oxygen transport from arterioles to tissue for simplicity.

Adopting this assumption eliminates Process 5 and reduces the 3-compartment model to a 2-compartment model with *C*_*a*_ being a constant; the oxygen transport model now becomes

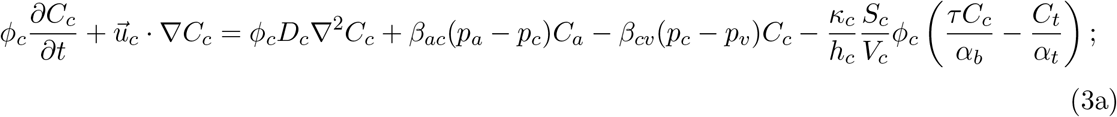

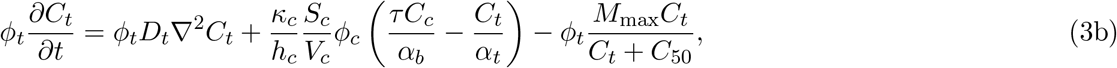

with homogenous NBC enforced on all the boundary surfaces of both compartments.

## 3 Parameter estimation

While the perfusion-related parameters are obtained through the perfusion model detailed in [19], values of the majority of the oxygen-related parameters are acquired from the literature. Table 2 summaries the chosen value for each parameter and their reference range. Both the coupling coefficients and the maximum consumption rate of oxygen have different values in the grey matter (GM) and the white matter (WM) due to the physiological differences between them [19, 37].

**Table 2:** Parameters values found in the literature. ^*^Reference ranges for *β*_*<sub>cv</sub>*_ are calculated as 3.5*β*_*<sub>ac</sub>*_ [19].

| Parameter | Chosen value | Reference range | Unit | Reference |
| --- | --- | --- | --- | --- |
| $\beta_{ac}^{GM}$ | 1.326 | $0.885 \pm 0.184$ | $\times 10^{-6} \text{ 1/(Pa}\cdot\text{s)}$ | [19–21] |
| $\beta_{ac}^{WM}$ | 0.522 | $0.253 \pm 0.062$ | $\times 10^{-6} \text{ 1/(Pa}\cdot\text{s)}$ | [19–21] |
| $\beta_{cv}^{GM}$ | 4.641 | $3.098 \pm 0.644^*$ | $\times 10^{-6} \text{ 1/(Pa}\cdot\text{s)}$ | [19–21] |
| $\beta_{cv}^{WM}$ | 1.828 | $0.886 \pm 0.217^*$ | $\times 10^{-6} \text{ 1/(Pa}\cdot\text{s)}$ | [19–21] |
| $C_a$ | 0.2 | $0.187 - 0.205$ | $\text{mm}^3 \text{O}_2/\text{mm}^3 \text{ blood}$ | see text |
| $D_c$ | 1.6 | $1.57 - 1.65$ | $\times 10^{-3} \text{ mm}^2/\text{s}$ | [38, 39] |
| $D_t$ | 1.8 | $1.2 - 2.0$ | $\times 10^{-5} \text{ mm}^2/\text{s}$ | [40–42] |
| $\kappa_c$ | 5.0 | $4.2 - 6.8$ | $\times 10^{-8} \text{ mm}^3 \text{O}_2/(\text{mmHg} \cdot \text{mm} \cdot \text{s})$ | [16, 41, 43, 44] |
| $h_c$ | 0.309 | $0.3 - 1$ | $\times 10^{-3} \text{ mm}$ | [44–47] |
| $\alpha_b$ | 3.11 | $2.86 - 3.11$ | $\times 10^{-5} \text{ mm}^3 \text{O}_2/(\text{mm}^3 \text{ blood} \cdot \text{mmHg})$ | [48–50] |
| $\alpha_t$ | 2.6 | $1.97 - 3.95$ | $\times 10^{-5} \text{ mm}^3 \text{O}_2/(\text{mm}^3 \text{ tissue} \cdot \text{mmHg})$ | [40–42] |
| $\tau$ | 0.01 | $0.008 - 0.015$ | - | [51] |
| $M_{\max}^{GM}$ | 7.77 | $5.05 - 10.33$ | $\times 10^{-4} \text{ mm}^3 \text{O}_2/(\text{mm}^3 \text{ tissue} \cdot \text{s})$ | [37, 42, 52, 53] |
| $M_{\max}^{WM}$ | 2.91 | $2.05 - 4.73$ | $\times 10^{-4} \text{ mm}^3 \text{O}_2/(\text{mm}^3 \text{ tissue} \cdot \text{s})$ | [37, 42, 52, 53] |
| $C_{50}$ | 2.6 | $0.078 - 10.4$ | $\times 10^{-5} \text{ mm}^3 \text{O}_2/(\text{mm}^3 \text{ tissue})$ | see text |

The value of *C*_*a*_ is calculated using the following relationship

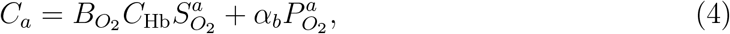

where 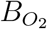 is the Hüfner constant representing the oxygen binding capacity of haemoglobin, which has a theoretical value of 1.39 [mL O_2_/g Hb] [54]; *C*_Hb_ is the haemoglobin concentration in normal blood that has a typical value of 0.15 [g Hb/mL blood] [55]. Oxygen saturation in the arterioles 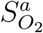 is related to 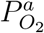 through the ODC, which can be approximated by the Hill equation

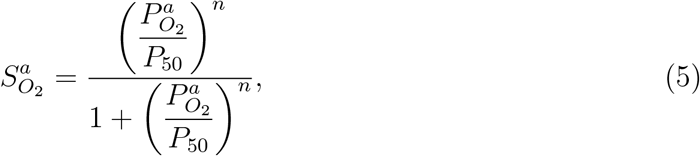

with *P*_50_ defined as the 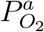 at 50% oxygen saturation with a value of 26.2 [mmHg] and *n* being the Hill coefficient with a value of 2.5 for normal human adults [56]. As values of 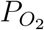 for cortical vessels are hard to measure and data on human are rare, we reference the measurements in animal cortex which suggest a 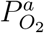 of 60–100 [mmHg] [34, 57–63]. Choosing a mean value of 80 [mmHg], *C*_*a*_ is estimated to be 0.2 [mm^3^ O_2_/mm^3^ blood] for a healthy individual.

The value of *C*_50_ is calculated through Henry’s law

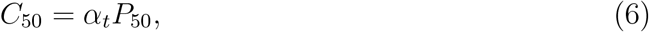

where *P*_50_ is the partial pressure of oxygen at half maximum consumption rate and has a range of 0.3–4 [mmHg] [64–67]. Adopting the chosen value of *α*_*t*_ as given in Table 2 together with *P*_50_ =1 [mmHg] results in *C*_50_ = 2.60 × 10^*−*5^ [mm^3^ O_2_/mm^3^ tissue]. Additionally, the reference range of *M*_max_ is simply assumed to be the cerebral metabolic rate of oxygen (CMRO_2_) reported in the literature. Values of CMRO_2_ provide a good approximation of *M*_max_ when *C*_*t*_ ≫ *C*_50_ which is the case under all healthy conditions.

The remaining parameters, *S*_*c*_/*V*_*c*_, *ϕ*_*c*_ and *ϕ*_*t*_, are the geometric parameters. They are estimated by employing synthetic geometries of the microvasculature that mirror the statistical properties of the human vasculature [12]. Although the arteriolar compartment is not included in the model, computation of its geometric parameters can be found in [31]. Capillary networks generated by El-Bouri and Payne [12] are utilised. Nine different cubical voxel sizes with their side length (*L*) ranging from 180 [*µ*m] to 625 [*µ*m] are selected. There are 500 networks for each voxel size, which enables the characterisation of the potential variability in the microvasculature. The diameters of the capillaries are assumed to be normally distributed with values of 6.3 ± 1.3 [*µ*m] [12].

Having the segment length (*l*) and the corresponding radius (*r*) of each vessel extracted for each voxel, the total capillary volume and surface area are calculated as

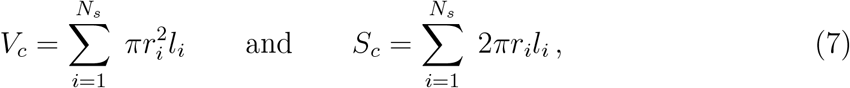

respectively, where *N*_*s*_ represents the total number of vessel segments within a given voxel. The voxel volume is calculated as *V*_*vox*_ = *L*^3^. It is then straightforward to evaluate *S*_*c*_/*V*_*c*_ and *ϕ*_*c*_ is computed as *V*_*c*_/*V*_*vox*_. In addition to *ϕ*_*c*_ and *S*_*c*_/*V*_*c*_, two more parameters are calculated for the purpose of validating against literature values; these parameters are the vessel volume (including blood) to vessel surface area ratio (*V*_*c*_/*S*_*c*_) and the vessel surface area to voxel volume ratio (*S*_*c*_/*V*_*vox*_).

Table 3 summarises the mean parameter values for each voxel size. To characterise the statistics of each voxel size, all parameters were tested for normal distribution using the Anderson-Darling test [68] with a significance level of 1%. All parameters failed to reject the null hypothesis except *ϕ*_*c*_ and *S*_*c*_/*V*_*vox*_ of the 180 *µ*m cube. The reason for this could be that the number of vessel segments presented in this cube size is relatively small giving rise to a discrete distribution of these two parameters.

**Table 3:** Mean parameter values and their standard deviations of each voxel size. ^*^indicates the parameters that rejected the null hypothesis.

| Voxel size [$\mu\text{m}$] | $\phi_c$ [%] | $\frac{S_c}{V_c} \left[ \frac{\text{mm}^2}{\text{mm}^3} \right]$ | $\frac{V_c}{S_c} [\mu\text{m}]$ | $\frac{S_c}{V_{vox}} \left[ \frac{\text{mm}^2}{\text{mm}^3} \right]$ |
| --- | --- | --- | --- | --- |
| 180 | $1.55 \pm 0.103^*$ | $590.65 \pm 9.647$ | $1.69 \pm 0.028$ | $9.18 \pm 0.543^*$ |
| 250 | $1.57 \pm 0.067$ | $593.57 \pm 7.383$ | $1.69 \pm 0.021$ | $9.34 \pm 0.336$ |
| 300 | $1.38 \pm 0.042$ | $600.57 \pm 5.711$ | $1.67 \pm 0.016$ | $8.30 \pm 0.217$ |
| 375 | $1.43 \pm 0.033$ | $607.11 \pm 5.010$ | $1.65 \pm 0.014$ | $8.70 \pm 0.160$ |
| 425 | $1.41 \pm 0.026$ | $610.44 \pm 4.041$ | $1.64 \pm 0.011$ | $8.60 \pm 0.129$ |
| 450 | $1.44 \pm 0.025$ | $610.47 \pm 3.979$ | $1.64 \pm 0.011$ | $8.79 \pm 0.120$ |
| 500 | $1.41 \pm 0.021$ | $615.58 \pm 3.282$ | $1.62 \pm 0.0087$ | $8.69 \pm 0.104$ |
| 550 | $1.45 \pm 0.018$ | $614.86 \pm 2.772$ | $1.63 \pm 0.0073$ | $8.90 \pm 0.085$ |
| 625 | $1.42 \pm 0.015$ | $616.38 \pm 2.250$ | $1.62 \pm 0.0059$ | $8.72 \pm 0.074$ |

To obtained the converged parameters, best-fit curves of the parameter means are identified using the following form:

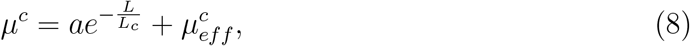

where *µ*^*c*^ is the mean of the parameters at every voxel size, *a* is a stretch coefficient, *L* is the voxel side length, *L*_*c*_ is a characteristic length and 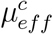 is the effective parameter.

The root mean square error (RMSE),

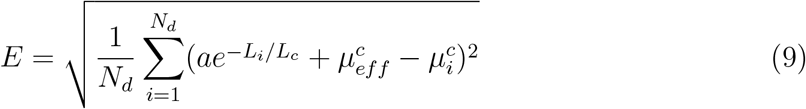

with *N*_*d*_ being the total number of data points, between the parameter mean and its best-fit curve is minimised and values of *a, L*_*c*_ and 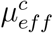 are found using the `fminsearch` function in MATLAB [69]. Values of the coefficients and the fitting errors are shown in Table 4.

**Table 4:** Best-fit curve coefficients of the capillary parameter means.

| | $a$ | $L_{ch} [\mu\text{m}]$ | $\mu_{eff}^c$ | %E |
| --- | --- | --- | --- | --- |
| $\phi_c$ | 0.98 % | 97.82 | 1.42 % | 2.75 |
| $\frac{S_c}{V_c}$ | $-67.32 \frac{\text{mm}^2}{\text{mm}^3}$ | 319.52 | $627.31 \frac{\text{mm}^2}{\text{mm}^3}$ | 0.25 |
| $\frac{V_c}{S_c}$ | 0.19 $\mu\text{m}$ | 304.03 | 1.59 $\mu\text{m}$ | 0.25 |
| $\frac{S_c}{V_{vox}}$ | $7.19 \frac{\text{mm}^2}{\text{mm}^3}$ | 69.69 | $8.70 \frac{\text{mm}^2}{\text{mm}^3}$ | 2.68 |

Data in Table 3 are visualised in Figures 2a–2d together with the best-fit curves of the parameter means illustrated as blue lines. It can be seen in Figures 2a and 2d that both *ϕ*_*c*_ and *S*_*c*_/*V*_*vox*_ of the 250 *µ*m cube and the 300 *µ*m cube appear to deviate the most from their best-fit curves. For the 250 *µ*m cube, the combination of having a relatively high mean segment density (7991/mm^3^ compared with a range of 7685–8272/mm^3^ [12]) and a large mean segment length (57.2 [*µ*m] compared with a range of 53.5–57.2 [*µ*m] [12]) is responsible for the high mean values of *ϕ*_*c*_ and *S*_*c*_/*V*_*vox*_. On the other hand, the segment density and the mean segment length of the 300 *µ*m cube are 7685/mm^3^ (range 7685– 8272/mm^3^) and 53.5 [*µ*m] (range 53.5–57.2 [*µ*m]), respectively [12], which are at the low end of their respective range. This results in smaller mean values of *ϕ*_*c*_ and *S*_*c*_/*V*_*vox*_.

**Figure 2.**
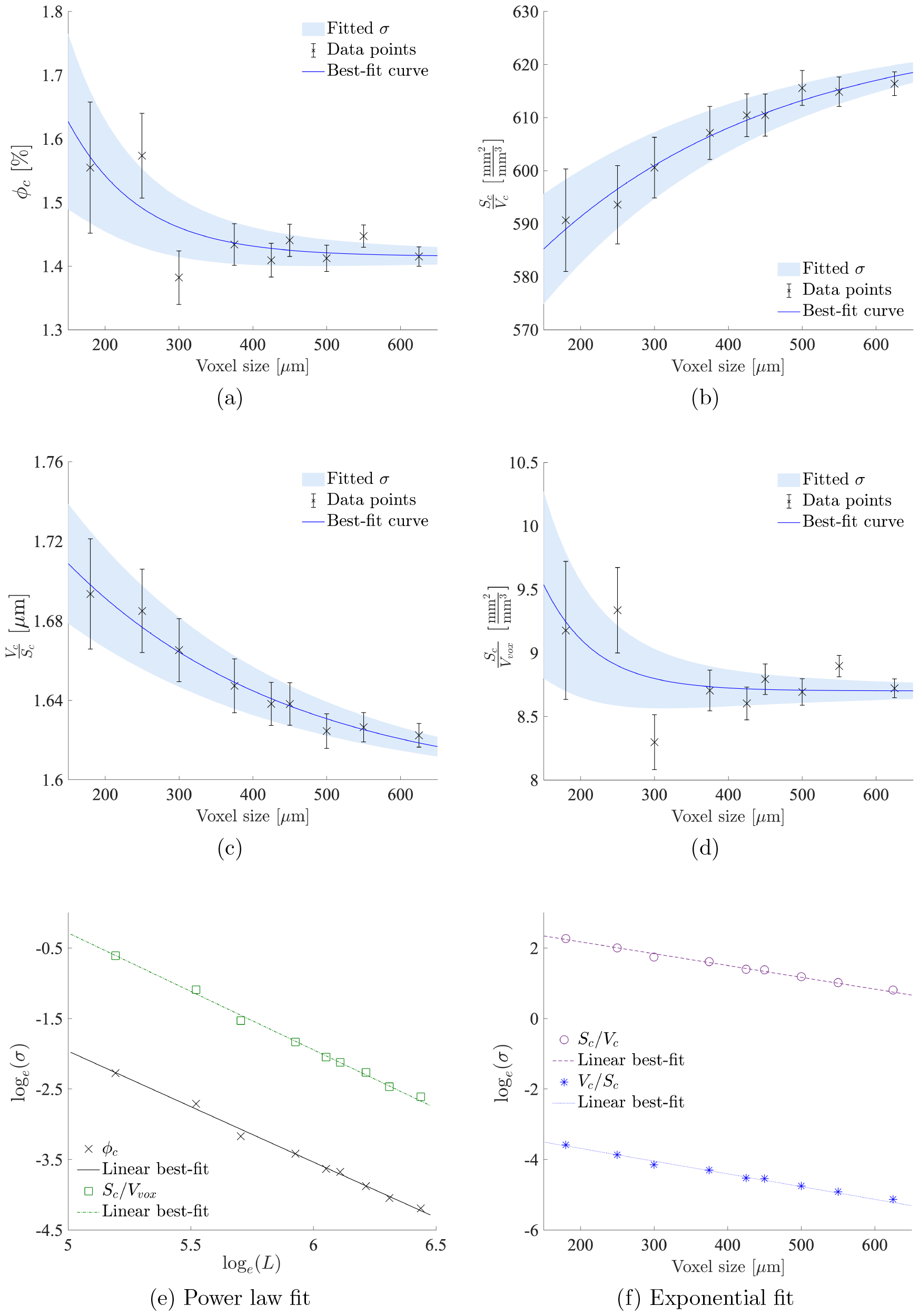
Error bar plots of means and standard deviations (data points), best-fit curves (blue line) and fitted standard deviations (light blue shadow) of the capillary parameters against voxel size (a–d); capillary standard deviations and their best-fit lines with (e) log-log scaling and (f) log-linear scaling.

Additionally, for the 550 *µ*m cube, the mean segment density is at the high end of the range with a value of 8272/mm^3^ (range 7685–8272/mm^3^), while its mean segment length sits near the middle of the range with a value of 54.9 [*µ*m] (range 53.5–57.2 [*µ*m]) [12]. Therefore, the mean values of *ϕ*_*c*_ and *S*_*c*_/*V*_*vox*_ of this voxel size deviate slightly from the best-fit curves.

Regardless of these variations, both *ϕ*_*c*_ and *S*_*c*_/*V*_*vox*_ converged with acceptable root mean square errors. In contrast, the best-fit curves of *S*_*c*_/*V*_*c*_ and *V*_*c*_/*S*_*c*_ did not reach convergence despite having very small fitting errors. However, 625 *µ*m remained as the upper limit of the cube size used in this work as the difference (around 1.84%) between the effective values and the means of the three largest cubes can be considered small.

The standard deviations (*σ*) of all parameters become smaller as the voxel size increases, indicating a reduction in morphological variability among individual capillary networks. Linear reduction in *σ*(*ϕ*_*c*_) and *σ*(*S*_*c*_/*V*_*vox*_) with increasing voxel size is shown when plotted on a log-log scale (Figure 2e). The same trend can be observed for *σ*(*S*_*c*_/*V*_*c*_) and *σ*(*V*_*c*_/*S*_*c*_) on a log-linear scale (Figure 2f). Because of these observations, power functions of the form

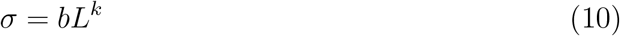

were chosen to represent the standard deviations of *ϕ*_*c*_ and *S*_*c*_/*V*_*vox*_, and exponential functions of the form

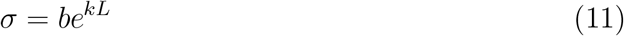

were chosen for the rest. The best-fit lines were found using the Curve Fitting Tool in MATLAB [69]. RMSE was used for evaluating the goodness of fit; the closer the value is to 0, the better the fit is. The coefficient values and the RMSEs are reported in Table 5. Linear representations of the fitted lines are shown in Figures 2e and 2f and the fitted *σ* for each parameter are shown as blue shades in Figures 2a–2d.

**Table 5:** Best-fit line coefficients of the standard deviation of capillary parameters. ^*^using power fit with Equation (10); ^*†*^using exponential fit with Equation (11).

| | $\phi_c^*$ | $\frac{S_c^\dagger}{V_c}$ | $\frac{V_c^\dagger}{S_c}$ | $\frac{S_c^*}{V_{vox}}$ |
| --- | --- | --- | --- | --- |
| $b$ | 365.9 | 17.17 | 0.0517 | 2976 |
| $k$ | -1.573 | -0.00335 | -0.00361 | -1.657 |
| RMSE | 0.0025 | 0.26 | 0.00077 | 0.010 |

The geometric parameters extracted from statistically accurate capillary beds are then validated against the ones obtained from physiologically accurate networks (Table 6). Values provided by Cassot *et al*. [70] and Lauwers *et al*. [71] were acquired from sections of a human brain; whereas, Risser *et al*. [72] and Weber *et al*. [73] quantified these values using monkeys. Only the study by Weber *et al*. [73] provided capillary statistics for all three parameters. For the other studies, only *ϕ* was quantified for the capillary vessels alone. As shown in Table 6, the statistically obtained *ϕ*_*c*_ agrees well with the physiological values.

**Table 6:**
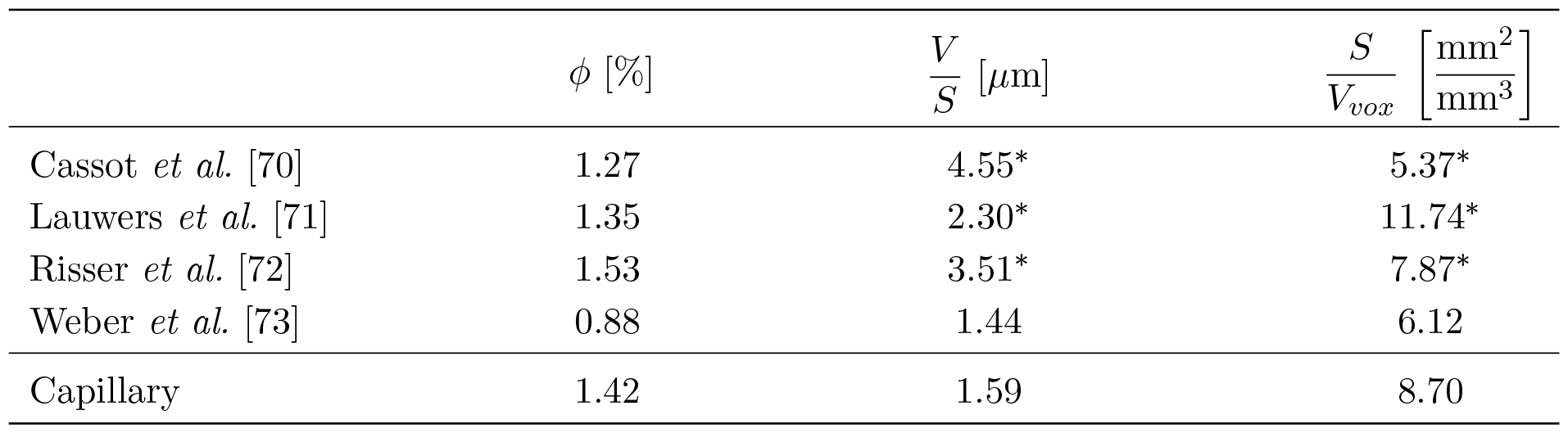
Comparison between the statistically computed capillary oxygen parameters and the physiological values in the literature. The cut-off diameters for capillaries are 9 [*µ*m], 10 [*µ*m], 11.2 [*µ*m] and 8 [*µ*m] for Cassot *et al*. [70], Lauwers *et al*. [71], Risser *et al*. [72] and Weber *et al*. [73], respectively. ^*^ includes non-capillaries vessels.

| | $\phi$ [%] | $\frac{V}{S}$ [$\mu\text{m}$] | $\frac{S}{V_{vox}}$ $\left[\frac{\text{mm}^2}{\text{mm}^3}\right]$ |
| --- | --- | --- | --- |
| Cassot *et al.* [70] | 1.27 | 4.55* | 5.37* |
| Lauwers *et al.* [71] | 1.35 | 2.30* | 11.74* |
| Risser *et al.* [72] | 1.53 | 3.51* | 7.87* |
| Weber *et al.* [73] | 0.88 | 1.44 | 6.12 |
| Capillary | 1.42 | 1.59 | 8.70 |

Parameters *V*_*c*_/*S*_*c*_ and *S*_*c*_/*V*_*vox*_ are slightly higher than their corresponding capillary-only measurements but are within the overall range. However, Weber *et al*. [73] used a lower value for the cut-off diameter which is reflected in its lower volume fraction; values of *V*_*c*_/*S*_*c*_ and *S*_*c*_/*V*_*vox*_ should be higher in networks with a larger volume fraction. It is also worth noting that there is a rather large variation between the values of *S*/*V*_*vox*_ even when the samples used are obtained from the same region [70, 71]. As tissue is the sole occupant of the remaining volume in this two-compartment model, *ϕ*_*t*_ is then calculated as 1 − *ϕ*_*c*_, giving a value of 98.52%.

Finally, in addition to the REV identified base on the permeabilities of the capillary networks [12], an REV can also be calculated now based on the oxygen parameters by comparing the voxel means and the effective values. An acceptable error margin is set to decide the minimum REV. Figure 3 shows the percentage errors, 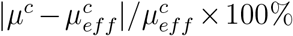, of each parameter for all 9 voxel sizes. In general, the error of all parameters reduces with increased voxel size as expected. In accordance with [12], the error tolerance is chosen to be 4.5% and the 375 *µ*m cube can thus be identified as the REV for this network based on the oxygen parameters. This REV is the same as the value suggested by perfusion [12], which is well below typical MRI resolution and is thus recommended for future studies on oxygen transport in discrete capillary networks.

**Figure 3.**
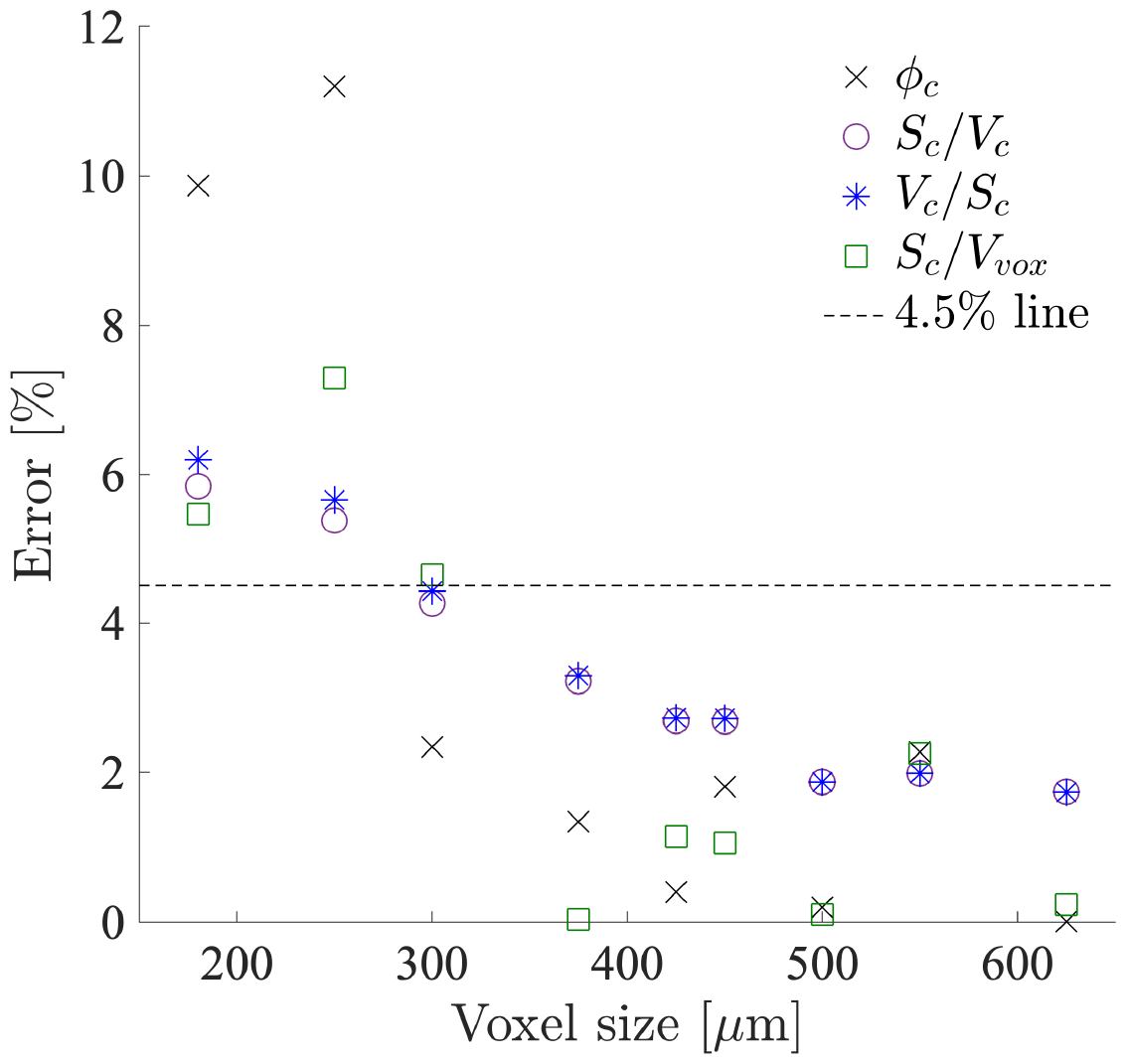
Percentage errors of the capillary voxel parameter means.

## 4 Model implementation

Since the oxygen transport model makes use of the results from the perfusion model as inputs, the same numerical method, the finite element method (FEM), is also employed here. A variational form of Equation (3) is firstly derived and then discretised by the continuous Bubnov-Galerkin method [74]. FEniCS [75], an open source finite element library, is employed here to obtain the numerical solutions.

Oxygen concentration in each compartment is discretised by first-order continuous Galerkin elements. Second-order continuous Galerkin elements are assigned to *p*_*a*_, *p*_*c*_ and 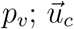 is subsequently represented by first-order continuous Galerkin elements and computed as

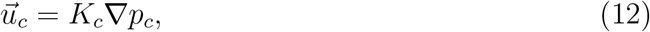

where *K*_*c*_ is the permeability of the capillary compartment with a value of 4.28×10^*−*7^ [mm^3^·s/g] [12, 19]. This combination of element choices has been shown to reduce the mass conservation error in the perfusion model [20]. Lastly both the coupling coefficients and the maximum consumption rate of oxygen are stored in discontinuous piecewise constant scalar function spaces.

For simplicity, we consider the steady state conditions which eliminates the temporal terms. The governing equations are first solved on a 1D tissue column to investigate the model behaviour and its sensitivity towards the parameter values. The 1D geometry is chosen for its low computational cost; it has been shown with the perfusion model that the 1D simulations are about 3000 times faster compared to the 3D brain simulations [20]. The model is then implemented within a patient-specific brain mesh detailed in [19]. This brain mesh consists of approximately 1 million tetrahedral elements, two computational subdomains (grey matter, Ω_GM_, and white matter, Ω_WM_), and three bounding surfaces (cortical surface, inner surface of the ventricles and a transverse cut of the brainstem).

### 4.1 1D tissue column

The 1D tissue column contains both grey and white matter, and all the model parameters are identical to those identified for the 3D brain model. The length of this tissue column was optimised by Józsa *et al*. [20], which has a value of 21.54 [mm]. The equivalent lengths of the grey and white matters are 13.55 [mm] and 7.99 [mm], respectively, which ensure that the volume-averaged perfusion values of the 1D model are the same as that of the 3D model. An interval mesh with 1000 elements is used for the 1D simulations.

Figure 4 shows the simulation results of the oxygen transport model with two sets of metabolic rates. The solid black lines denote results using the oxygen consumption rate listed in Mintun *et al*. [42] where the grey and white matters have CMRO_2_ values of 8.20 × 10^*−*4^ and 2.05×10^*−*4^ [mm^3^ O_2_/(mm^3^ tissue · s)], respectively. The blue dash lines represent results using the mean metabolic rates taken from An *et al*. [52] with the grey and white matters having a CMRO_2_ value of 8.67×10^*−*4^ and 4.26×10^*−*4^ [mm^3^ O_2_/(mm^3^ tissue · s)], respectively.

**Figure 4.**
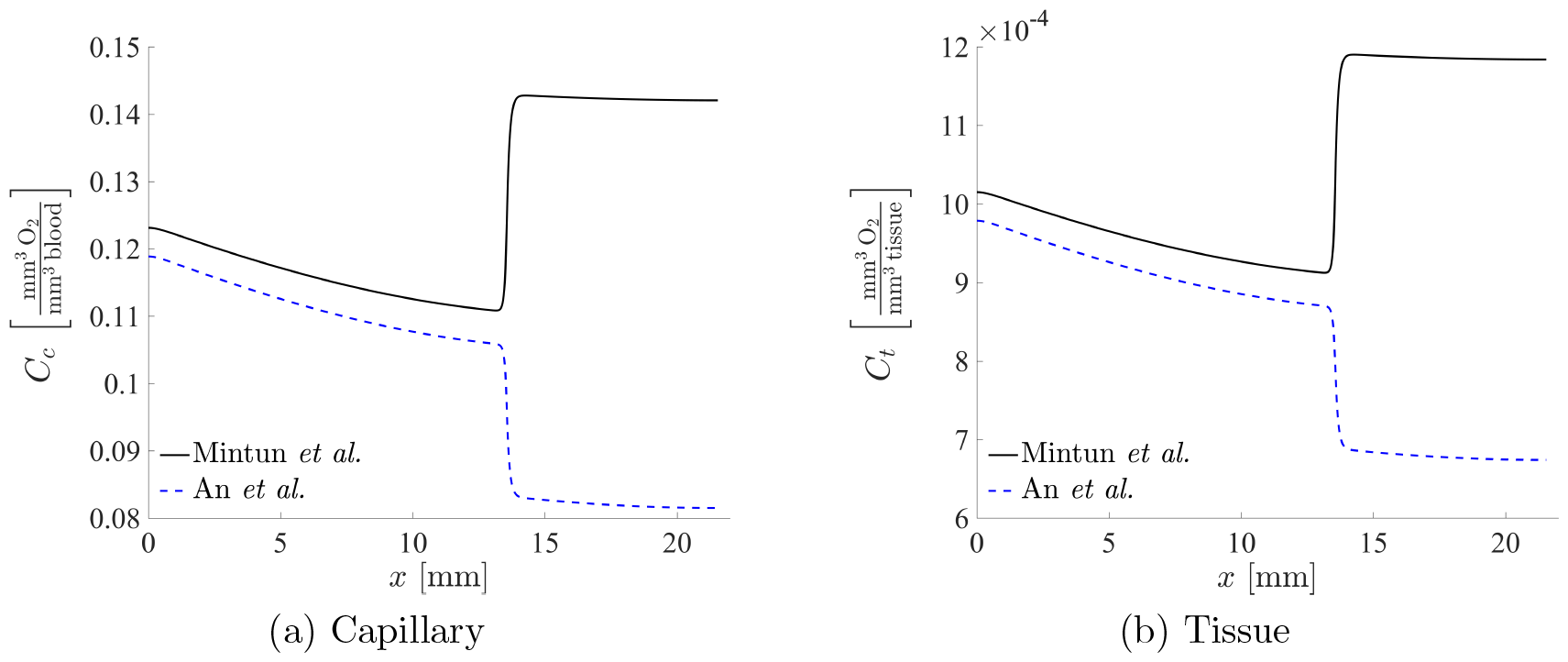
Simulation results of the oxygen transport model in a 1D tissue column with metabolic rate from Mintun *et al*. [42] (black solid line) and An *et al*. [52] (blue dash line). *x* = 0 [mm] indicates the cortical surface.

It can be seen that choosing different oxygen consumption rates leads to qualitatively different distributions of oxygen concentration in the grey and white matters. The source of oxygen for the capillary compartment is constant throughout the entire domain and the inter-compartment advection coefficients in the white matter drop to about a third of that in the grey matter. However, the grey-to-white matter metabolism ratio is 4:1, resulting in a higher oxygen level in the white matter. In contrast, An *et al*.’s [52] oxygen consumption rate in the white matter is about a half of that in the grey matter, leaving a lower oxygen level in the former.

As summarised in Table 2, the reported CMRO_2_ appears to have a large range; values for the grey and white matter range between 5.05–10.33×10^*−*4^ and 2.05–4.73×10^*−*4^ [mm^3^ O_2_/(mm^3^ tissue · s)], respectively. Since the choice of metabolic rate significantly affects the distribution of oxygen along the depth of the brain, the best way to find the most suitable value of *M*_max_ in this context thus appears to be through optimisation.

### 4.2 Parameter optimisation

To uniquely define the oxygen distribution, the values of 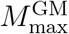 and 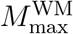 are optimised. The oxygen model is simplified into a set of algebraic equations through volume averaging by which to speed up the optimisation process. We consider the following assumptions. Firstly, 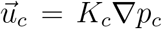 is negligible as *K*_*c*_ is relatively small [19]. Secondly, 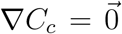 at the interface of grey and white matter. Lastly, the pressure within each compartment is constant which is shown to be well-approximated by their volume-averaged values [19].

Integrating Equation (3) over Ω_GM_, applying Gauss’s theorem and dividing by the grey matter volume then give the following algebraic equations for the grey matter:

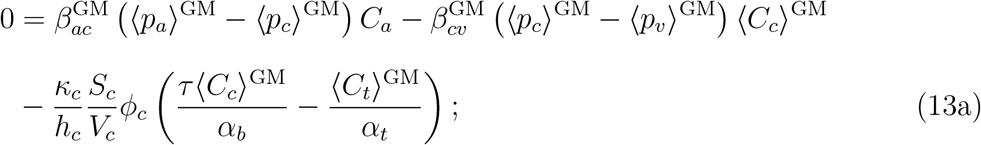

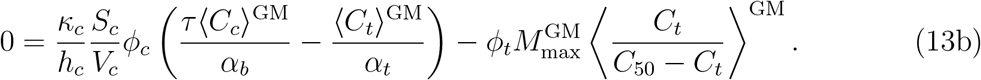

The angled brackets indicate volume-averaged values. The algebraic equation set for the white matter can be derived following the same process and the resultant equations are the same as that of the grey matter; therefore, the superscripts (GM or WM) are omitted unless necessary.

The challenge of this optimisation lies in finding a suitable criterion to construct the cost function. Although physiological values of cerebral partial pressure of oxygen have been reported, they vary among literature and few values have been reported specifically for the white matter. Oxygen extraction fraction (OEF), on the other hand, not only is relatively consistent across literature (OEF∈[0.3, 0.48]) but is almost identical between the grey and white matter [37]. Experimentally, OEF is measured via PET [3, 37, 76, 77] or, in recent years, via MRI [5, 78, 79] and is defined as

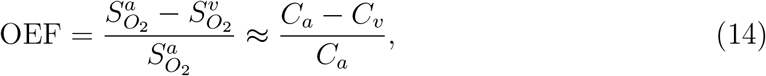

where 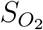 is oxygen saturation. However, the current oxygen model does not allow for the determination of the venular oxygen concentration (*C*_*v*_) due to the lack of venular compartment. To relate the algebraic equations of the oxygen model to the chosen optimisation criterion, we then consider the relationship between OEF and CMRO_2_, i.e.,

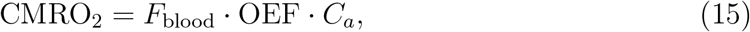

where *F*_blood_ is perfusion. According to [19], perfusion and the inter-compartment advection coefficients have the following relationship:

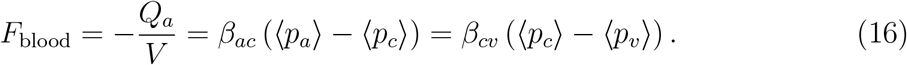

Now, substituting Equation (13a) and Equation (16) into Equation (13b) and rearranging the resultant equation lead to the following relationship

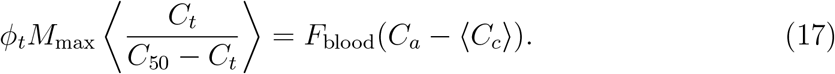

It can be seen that the left hand side represents CMRO_2_ of the system and the second term on the right hand side is equivalent to OEF · *C*_*a*_ from a volume-averaged perspective with the assumption that ⟨*C*_*v*_⟩ = ⟨*C*_*c*_⟩. Equation (17) is then used to optimise *M*_max_.

A value of 0.4 [37] is chosen as the value of OEF for both the grey and the white matter. The optimisation problem hence becomes finding *M*_max_ values that satisfy ⟨*C*_*c*_⟩ = 0.6*C*_*a*_ as inferred by Equations (15) and (17). Therefore, the cost function (*J*) to be minimised is

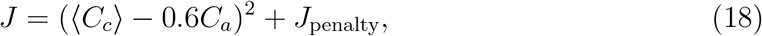

where *J*_penalty_ is a step function and follows

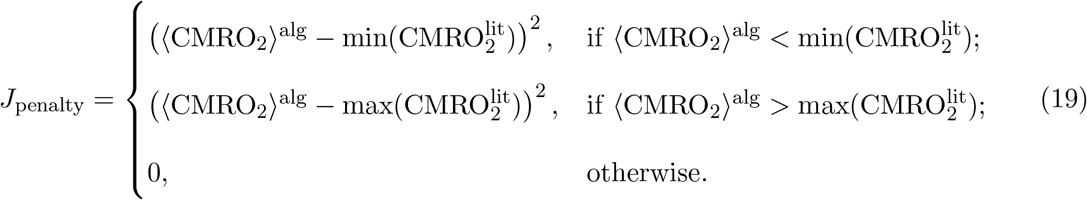

⟨CMRO_2_⟩^alg^ is the volume-averaged CMRO_2_ value computed as the left-hand-side term of Equation (17); 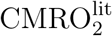 is the physiological values from the literature with its range listed in Table 2. The addition of the penalty term aims to restrict the metabolism of the numerical solution to be within the physiological range. Additionally, the bounding values of *M*_max_ are chosen to be the same as those for CMRO_2_.

The optimisation problem can now be phrased as

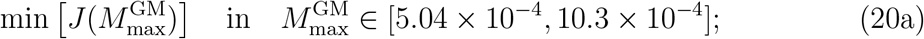

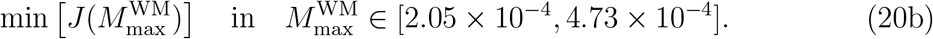

Note that optimisation of *M*_max_ for grey and white matter can be performed independently because *C*_*a*_ does not vary spatially and no-flow is assumed at the grey-white matter interface. To find the global minimum of *J*, two optimisation methods (Broy-den–Fletcher–Goldfarb–Shanno (BFGS) [80] and the Nelder-Mead [81]) are employed with the utilisation of Mathematica [82]. Both optimisation algorithms work with bounded optimisation; while BFGS is gradient based, Nelder-Mead does not require the computation of the cost function’s gradient.

Both optimisation algorithms are run three times and each with an initial guess generated randomly within the bounds, which ensures that the optimised *M*_max_ is independent from the choice of algorithms and the initial values. All runs converge rapidly towards the same *M*_max_ values reaching *J* < 10^*−*4^ in 30 iterations; convergence plots of *M*_max_ and *J* of the grey and white matter from a typical run are shown in Figure 5. The optimised parameter values are 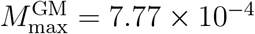 and 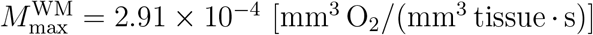.

**Figure 5.**
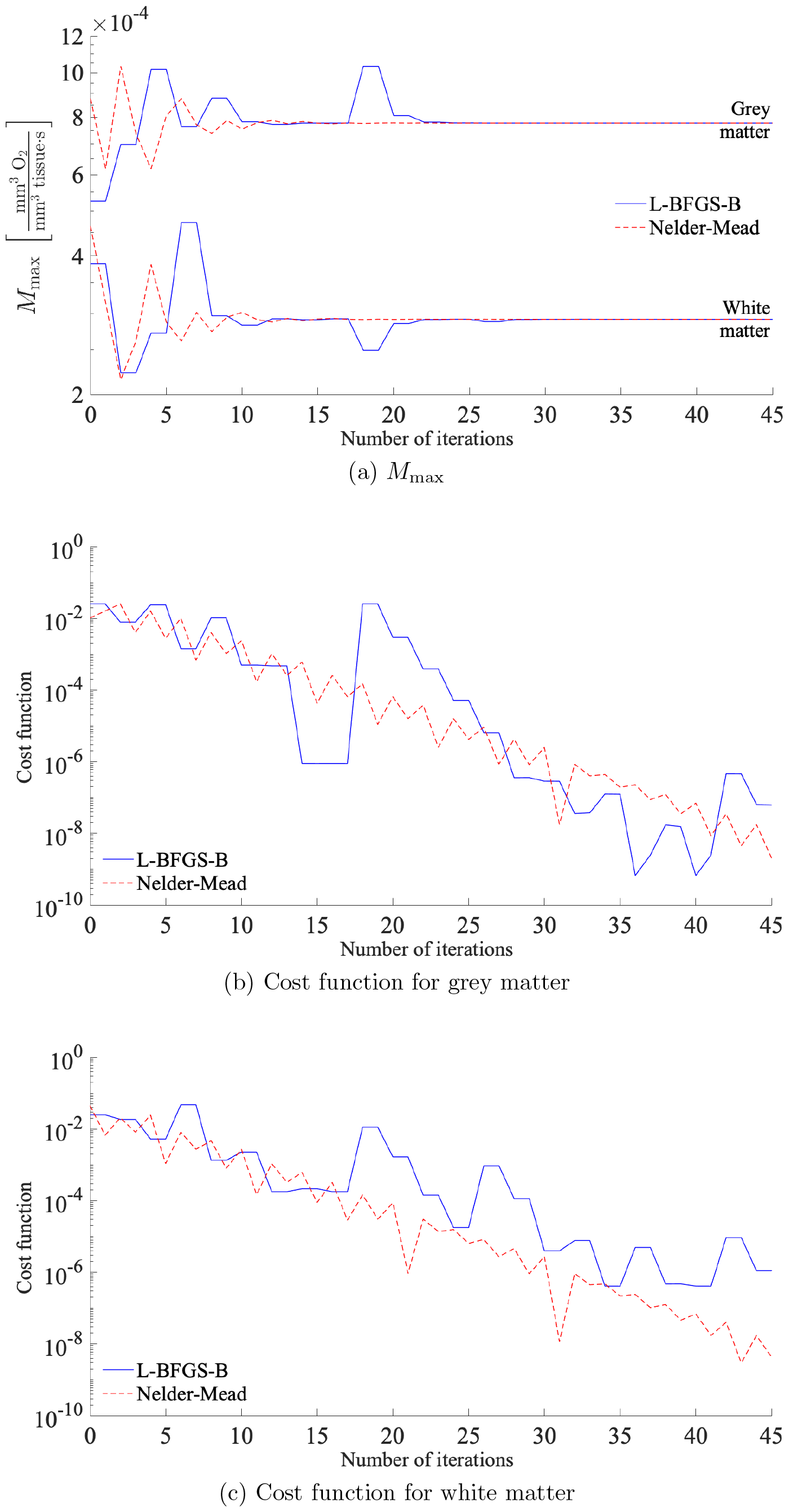
Convergence plots of *M*_max_ and cost function (*J*) of the grey and white matter from a typical optimisation.

## 5 Results

In this section, results of the 1D simulation using the optimised maximum consumption rates of oxygen are presented first followed by results of the 3D simulation within a patient-specific brain mesh. The results are examined quantitatively by calculating the volume-averaged values of the numerical solutions for each compartment and for each subdomain, i.e., the grey and white matters. Volume-averaged *C*_*i*_ and CMRO_2_ are calculated as

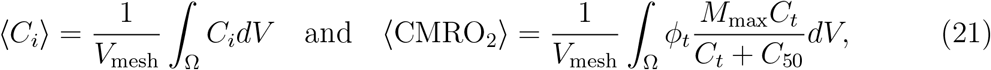

respectively, with *V*_mesh_ being the total volume of the domain Ω. To be consistent with the model, oxygen partial pressures are evaluated as

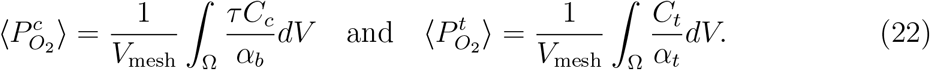

Finally, a sensitivity analysis is performed to quantify the uncertainties of the values employed by the model.

### 5.1 1D tissue column

Figure 6 shows the results of the oxygen transport model in a 1D tissue column with the optimised *M*_max_. Calculation of the OEF follows Equation (14) with the assumption that *C*_*c*_ = *C*_*v*_ under steady state. It can be seen from Figure 6a that the value of the OEF is maintained around the optimisation target with a less than 9% variation throughout the depth.

**Figure 6.**
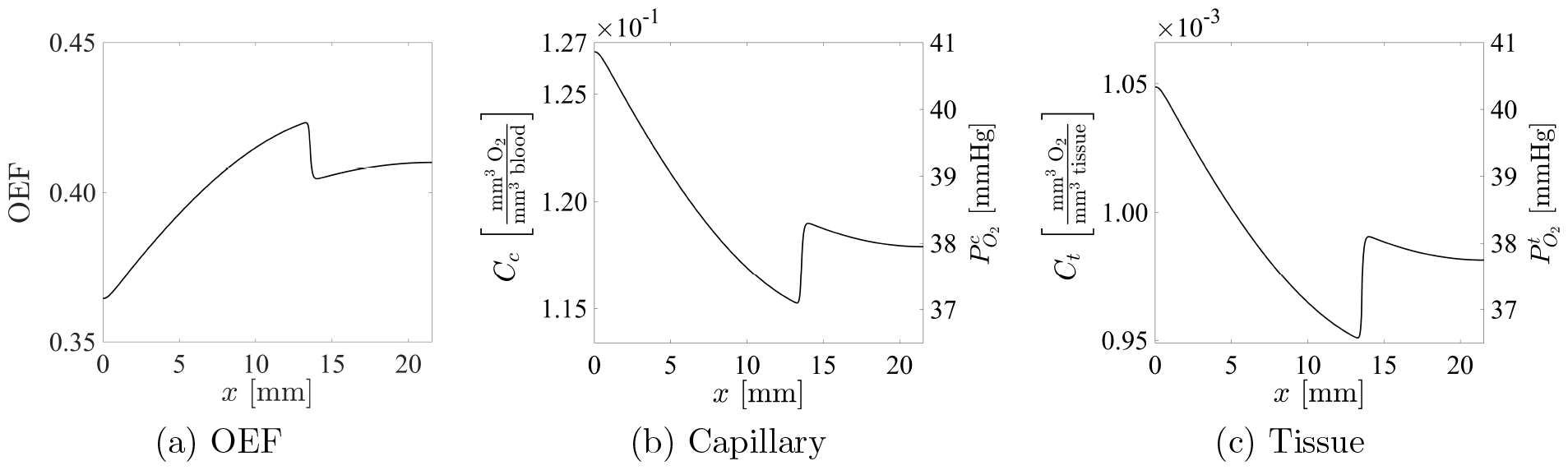
Simulation results of the 2-compartment oxygen model in a 1D tissue column with optimised metabolic rates.

Volume-averaged oxygen levels and consumption rates of both the whole domain and the subdomains are summarised in Table 7. Capillary and tissue 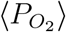 are of the same level as expected under steady state and they are within the physiological range of 28–60 [mmHg] [58, 59, 62, 63] and 22–45 [mmHg] [83–85], respectively. The volume-averaged CMRO_2_ is also in the physiological range.

**Table 7:**
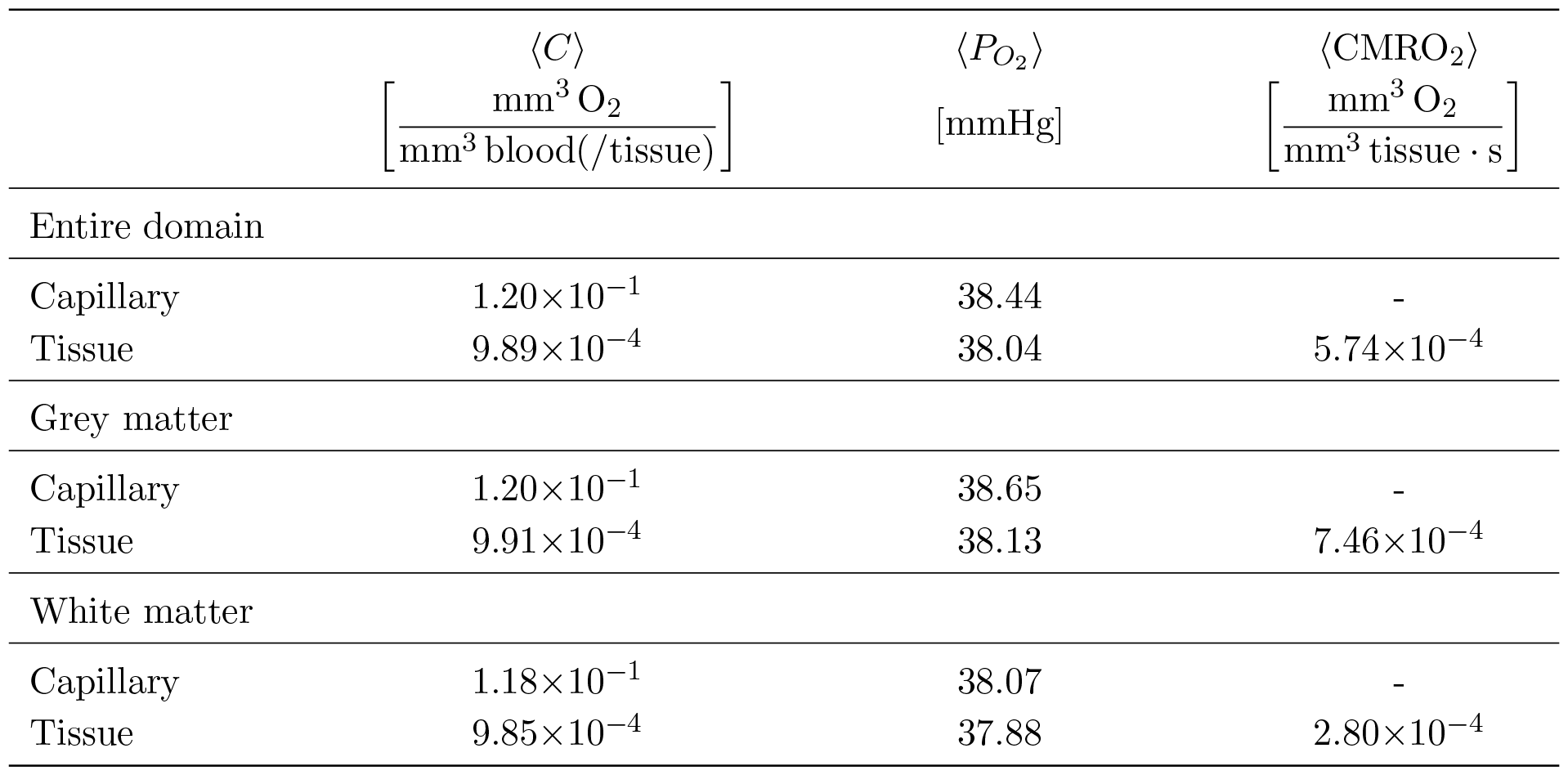
Volume-averaged values of oxygen concentration, partial pressure and CMRO_2_ of the oxygen transport model in a 1D tissue column with optimised metabolic rates.

| | $\langle C \rangle$<br>$\left[ \frac{\text{mm}^3 \text{ O}_2}{\text{mm}^3 \text{ blood(/tissue)}} \right]$ | $\langle P_{\text{O}_2} \rangle$<br>[mmHg] | $\langle \text{CMRO}_2 \rangle$<br>$\left[ \frac{\text{mm}^3 \text{ O}_2}{\text{mm}^3 \text{ tissue} \cdot \text{s}} \right]$ |
| --- | --- | --- | --- |
| Entire domain |  |  |  |
| Capillary | $1.20 \times 10^{-1}$ | 38.44 | - |
| Tissue | $9.89 \times 10^{-4}$ | 38.04 | $5.74 \times 10^{-4}$ |
| Grey matter |  |  |  |
| Capillary | $1.20 \times 10^{-1}$ | 38.65 | - |
| Tissue | $9.91 \times 10^{-4}$ | 38.13 | $7.46 \times 10^{-4}$ |
| White matter |  |  |  |
| Capillary | $1.18 \times 10^{-1}$ | 38.07 | - |
| Tissue | $9.85 \times 10^{-4}$ | 37.88 | $2.80 \times 10^{-4}$ |

Unlike the oxygen distributions shown in Figure 4, the optimised *M*_max_ leads to a decrease in oxygen level from the cortical surface towards the grey-white matter interface, reaching its lowest values, followed by an increase and then a gradual reduction in oxygen level. This increase in oxygenation at the grey-white matter interface seems counter-intuitive; however, it has not been determined experimentally how oxygen is distributed through the depth of the brain. Measurements regarding oxygen partial pressure versus brain depth are very scarce and both increasing and decreasing in tissue oxygenation with depth have been observed in human and animal studies [85–88].

Interestingly, experiments on rats conducted by Feng *et al*. [89] showed that 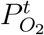 decreased with depth in the cortex and increased around the region of white matter. Differences of 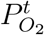 at 1000 [*µ*m], 2000 [*µ*m] and 3000 [*µ*m] underwent statistical evaluation; meanwhile, histological test confirmed that these depths were located within grey matter, white matter and hippocampal grey matter. It was found that 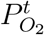 measured at 1000 [*µ*m] was significantly lower than that at 2000 [*µ*m].

A qualitative comparison between Feng *et al*.’s measurements and the simulation results is carried out by normalising each dataset through min-max scaling. Data of the mean 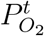 as a function of depth for closed skull rats are extracted from Fig. 2 in [89]. The original bounding values of the mean 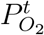 and depths are approximately (5; 44) [mmHg] and (0; 3000) [*µ*m], respectively. It can be seen in Figure 7 that, despite anatomical differences between the rat and the human brain, the 1D simulation result shows a very good qualitative agreement with the experimental data. The higher oxygen level in the white matter is mainly due to the higher oxygen content in the blood as *C*_*a*_ is constant over the domain.

**Figure 7.**
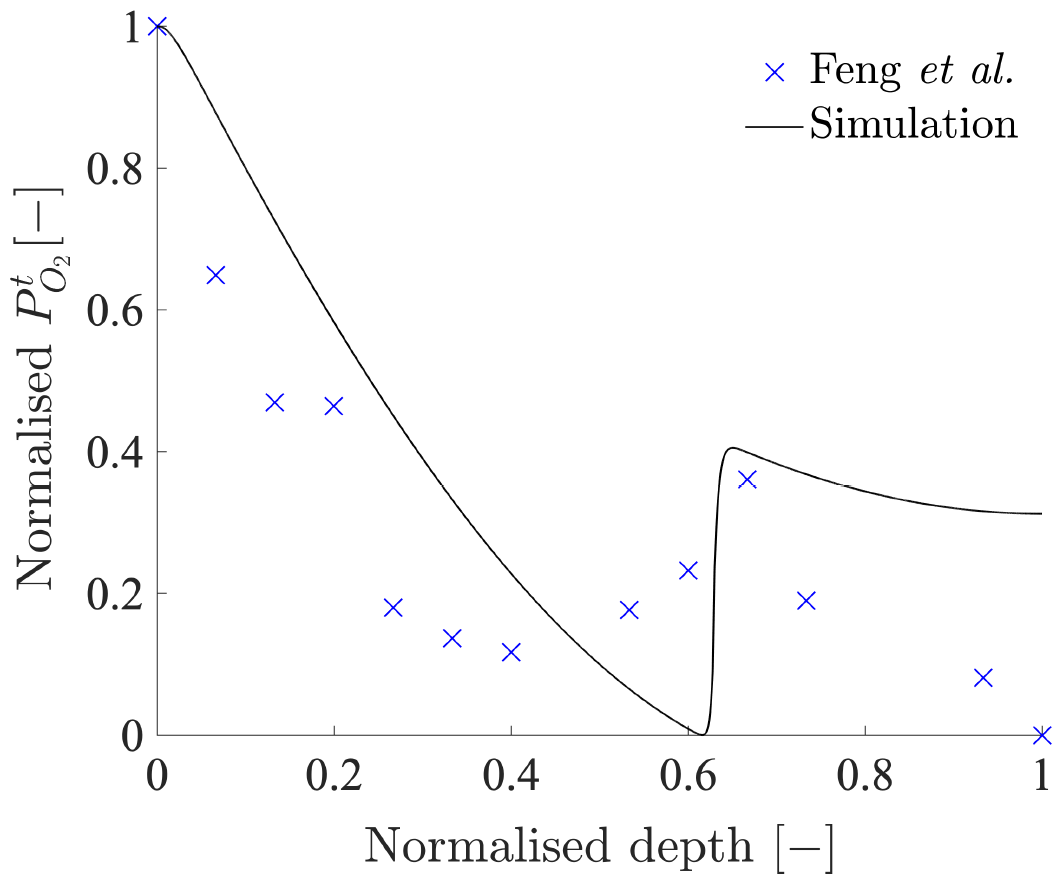
Comparison between mean 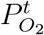 of closed skull rats [89] and simulated 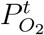 throughout depth. Both the experimental measurements and the simulation result have undergone independent min-max normalisation.

### 5.2 3D human brain

Simulation of a health scenario is next performed on the 3D brain mesh. The simulation is carried out on a high performance computer using 10 cores of an Intel Xeon Platinum 8268 processor with 80 GB RAM, which takes approximately 35 minutes. Volume-averaged values of the 3D results are summarised in Table 8.

**Table 8:**
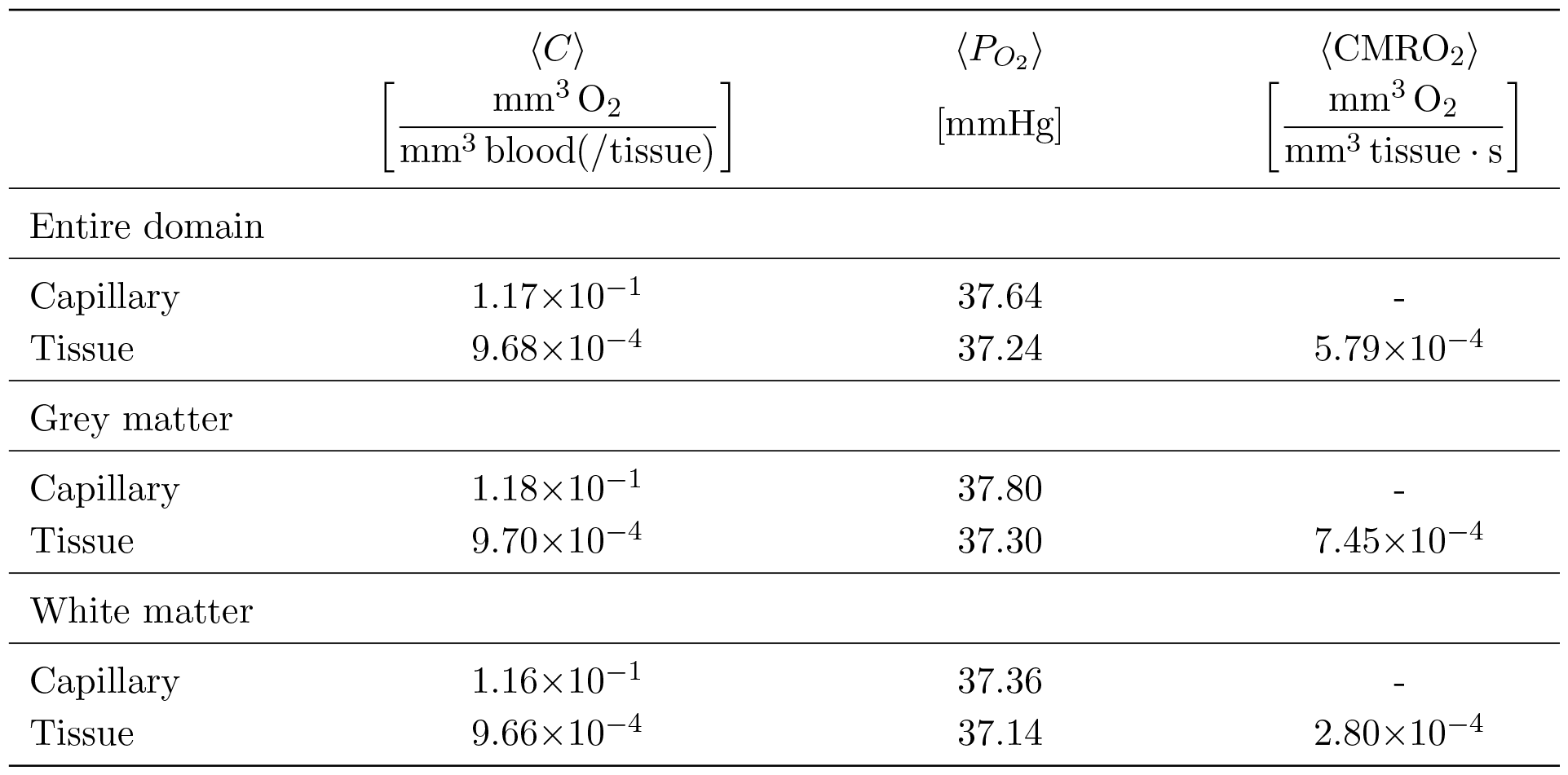
Volume-averaged values of oxygen concentration, partial pressure and CMRO_2_ of the oxygen transport model in a 3D human brain mesh.

| | $\langle C \rangle$<br>$\left[ \frac{\text{mm}^3 \text{O}_2}{\text{mm}^3 \text{blood}(/ \text{tissue})} \right]$ | $\langle P_{O_2} \rangle$<br>[mmHg] | $\langle \text{CMRO}_2 \rangle$<br>$\left[ \frac{\text{mm}^3 \text{O}_2}{\text{mm}^3 \text{tissue} \cdot \text{s}} \right]$ |
| --- | --- | --- | --- |
| Entire domain |  |  |  |
| Capillary | $1.17 \times 10^{-1}$ | 37.64 | - |
| Tissue | $9.68 \times 10^{-4}$ | 37.24 | $5.79 \times 10^{-4}$ |
| Grey matter |  |  |  |
| Capillary | $1.18 \times 10^{-1}$ | 37.80 | - |
| Tissue | $9.70 \times 10^{-4}$ | 37.30 | $7.45 \times 10^{-4}$ |
| White matter |  |  |  |
| Capillary | $1.16 \times 10^{-1}$ | 37.36 | - |
| Tissue | $9.66 \times 10^{-4}$ | 37.14 | $2.80 \times 10^{-4}$ |

The volume-averaged oxygen concentration and partial pressure in both compartments are slightly lower than that of the 1D model. This leads to a slightly higher volume-averaged OEF with a value of 0.41. Although ⟨CMRO_2_⟩ of the 3D model is very similar to that of the 1D model in the grey and white matters, the overall ⟨CMRO_2_⟩ of the 3D model is slightly higher.

The difference between the 1D and 3D results are partially attributed to the fact that the chosen FEM is not conservative. Reducing the element size will reduce numerical errors in the 3D results; however, it will also significantly increase the computational cost. Another reason lies in that the 1D model does not take into account the curvature effects presented in the 3D brain mesh. Additionally, the 3D model and 1D model have different ‘volume’ ratio between the grey and white matters with grey matter occupying a greater proportion in the 3D model. This contributes to a higher overall *M*_max_, resulting in a higher ⟨CMRO_2_⟩ in the 3D case.

Spatial distribution of oxygen in the 3D brain are shown on representative slices in Figures 8a and 8b. The 3D results generally follow the 1D trend; oxygen level decreases from the cortical surface towards the deeper cerebral layers with an increase in its values at the grey-white matter interface. Despite having similar volume-averaged values, oxygen distribution in the 3D brain shows a much larger variation throughout the depth. The lowest oxygen level is located around the ventricles instead of the grey-white matter interface and its value is around 10 times smaller than the smallest 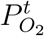 found in the 1D model.

**Figure 8.**
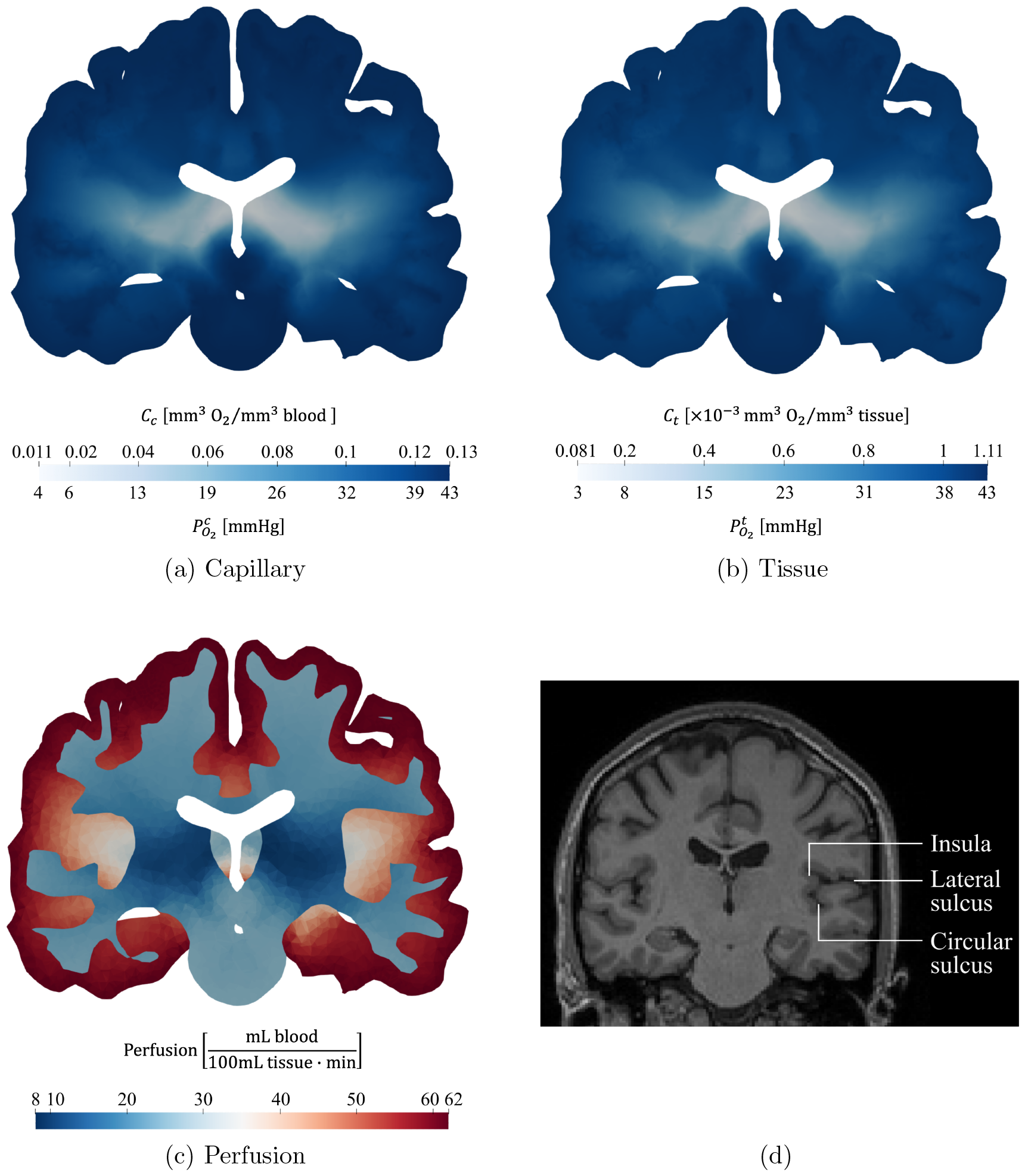
3D simulation results of (a) capillary oxygen distribution; (b) tissue oxygen distribution; (c) perfusion; and (d) a MRI image of a healthy subject IXI025 (adapted with annotation from [90] under license CC BY-SA 3.0 https://creativecommons.org/licenses/by-sa/3.0/).

The cause of this difference in oxygen level is rooted in the difference between the 1D and 3D perfusion values as a result of the anatomical inaccuracy of the current brain mesh [20,21]. Comparing the current mesh with Figure 8d, we can see that both the lateral and circular sulci are missing from the current mesh, especially in the right hemisphere. Although Figure 8d is not from the patient the mesh was constructed from, the lateral sulcus is the most prominent feature of the human brain. The space that should be filled with cerebrospinal fluid is occupied by the grey matter instead and the insula is unidentifiable from the mesh.

Even though the 1D and the 3D models have the same volume-averaged perfusion, they do not have the same distribution throughout the depth of the brain. The minimum perfusion value of the 3D model is 2.5 times smaller than that of the 1D model. This under perfused region is found in the white matter, near the intersection of the lateral and the third ventricles (see Figure 8c), which is located the furthest away from the cortical surface due to the missing sulci. This region corresponds well to the low oxygenation area.

### 5.3 Sensitivity analysis

One-at-a-time sensitivity analysis is performed using the 1D model and the parameters of interest are listed in Table 9. For a parameter Λ, its value is perturbed twice with values above (Λ^high^) and below (Λ^low^) the baseline value (Λ^base^). Both the baseline (chosen) value and the parameter range can be found in Table 2. Additionally, *ϕ*_*c*_ has a range of 0.83–2.36% [70–73] and a ±10% change is imposed on *S*_*c*_/*V*_*c*_. Percentage change of Λ relative to its baseline value is calculated as follow

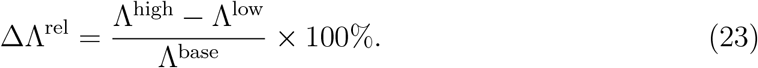

Sensitivity (ψ [%/%]) of the oxygen transport model towards a certain parameter is quantified as

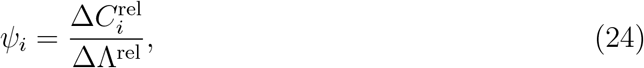

where 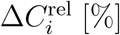 is the relative percentage change of oxygen concentration over the whole domain in compartment *i* and evaluated as

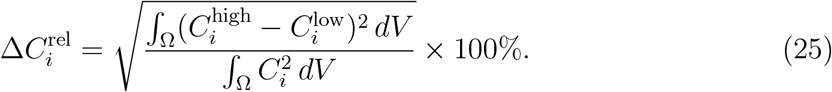

Here, *C*_*i*_ is the baseline oxygen concentration computed with Λ^base^; 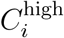 and 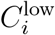 are the oxygen concentration acquired by employing Λ^high^ and Λ^low^, respectively.

**Table 9:** Relative percentage change of parameter values and the corresponding relative percentage change in capillary and tissue concentration.

| Parameter | $\Delta\Lambda^{\text{rel}}$ [%] | $\psi_c$ [%/%] | $\psi_t$ [%/%] |
| --- | --- | --- | --- |
| $D_c$ | 5.00 | $2.21 \times 10^{-4}$ | $2.23 \times 10^{-4}$ |
| $D_t$ | 44.4 | $2.31 \times 10^{-4}$ | $2.39 \times 10^{-4}$ |
| $\kappa_c$ | 52.0 | $1.86 \times 10^{-4}$ | $9.70 \times 10^{-3}$ |
| $h_c$ | 227 | $2.15 \times 10^{-4}$ | $1.11 \times 10^{-2}$ |
| $\alpha_b$ | 8.04 | $1.72 \times 10^{-2}$ | 1.08 |
| $\alpha_t$ | 76.2 | $2.12 \times 10^{-4}$ | 1.00 |
| $\tau$ | 70.0 | $1.44 \times 10^{-2}$ | 0.993 |
| $C_{50}$ | 397 | $1.58 \times 10^{-2}$ | $1.62 \times 10^{-2}$ |
| $\phi_c$ | 108 | $9.31 \times 10^{-3}$ | $2.07 \times 10^{-2}$ |
| $S_c/V_c$ | 20 | $2.14 \times 10^{-4}$ | $1.12 \times 10^{-2}$ |

Results of the sensitivity analysis is summarised in Table 9. All parameters have little effect on the capillary compartment; neither do most of the parameters on the tissue compartment. The model is most sensitive to the parameters in the inter-compartment diffusion terms. For every 1% relative change in *α*_*b*_, *α*_*t*_ or τ, there is approximately 1% change in the steady-state value of *C*_*t*_. This is because that oxygen transport from the capillary to tissue compartment is based on the 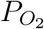 gradient which is related to oxygen concentration through oxygen solubilities.

As the chosen value of *α*_*b*_ is the upper bound in the reference range, 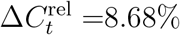 represents the maximum possible deviation from the model result for *α*_*b*_’s known physiological values. In contrast to the reference range of *α*_*b*_, *α*_*t*_ has a much wider range which gives raise to a larger uncertainty. However, Clark *et al*. [40] found that more than 50% of their measurements of *α*_*t*_ fell in the range of 2.37–2.89×10^*−*5^ [mm^3^ O_2_/(mm^3^ tissue · mmHg)], which reduces its 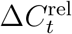 from 76.2% to 20%.

We now examine the effect of linearising the ODC has on the model results. Figure 9 illustrates the relationships between the blood oxygen concentration (*C*_*b*_) and the corresponding partial pressure of oxygen 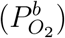. The linearisation with τ = 0.01 appears to overestimate 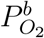 when 0.052 < *C*_*b*_ < 0.187 [mm^3^ O_2_/mm^3^ blood] and to underestimate 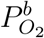 otherwise. While the range of *C*_*c*_ from the 1D model (narrow grey band) falls in the overestimated region, the range of *C*_*c*_ from the 3D model (wide blue band) overlaps with the overestimated and the lower portion of the underestimated regions. Although the linear approximation has little impact on values of *C*_*c*_ itself, it affects the corresponding 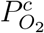 which in turn affects the oxygen level in tissue as shown in Table 9.

**Figure 9.**
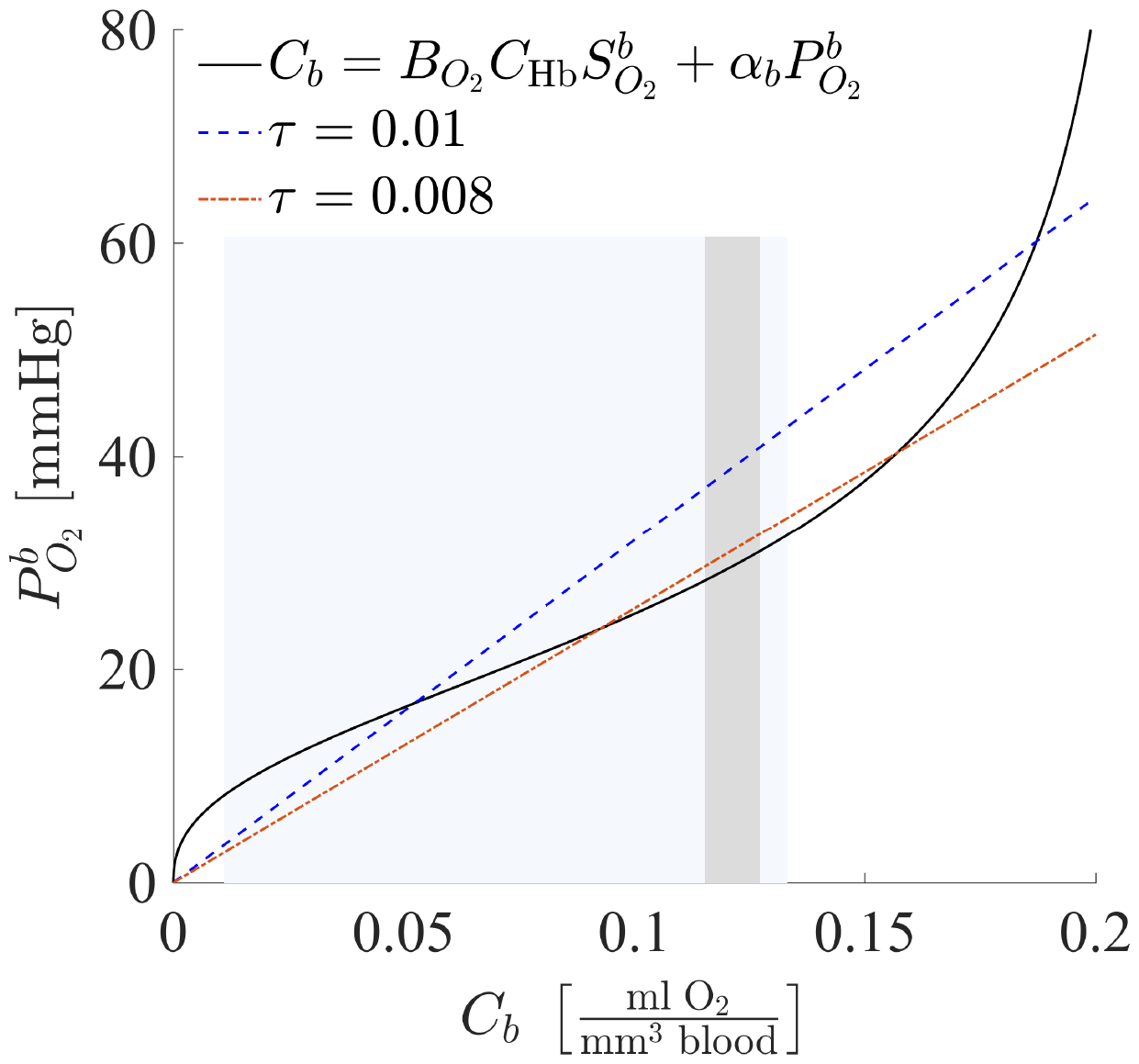
Relationship between 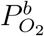 and *C*_*b*_: full relationship based on Hill equation approximated ODC (solid black line); linearised relationship 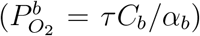 with τ = 0.01 (blue dashed line) and τ = 0.008 (orange dotted dash line). The narrow grey band and the wide blue band represent the range of *C*_*c*_ from the 1D and 3D models, respectively.

With τ = 0.008 — the lower bound of the reference range, a better overall approximation is achieved for both 1D and 3D models even though it does not give as good an estimation for the underestimated regions. For the 1D model, adopting τ = 0.08 gives 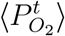 a value of 30 [mmHg] which is still within the physiological range. It is, therefore, possible that with an anatomically more accurate brain mesh, a linear approximation could also potentially provide a decent estimation for the 3D model under healthy scenarios.

## 6 Discussion

In this paper, we formulated a multi-scale, multi-compartment model describing organ-scale oxygen transport based on mass conservation. Geometric parameters of the model were quantified using statistically accurate microvascular networks. Volume fraction and vessel surface area to volume ratio of the capillary network converged to values of 1.42% and 627 [mm^2^/mm^3^], respectively, which are comparable to values obtained from human [70, 71] and monkey [72, 73] vascular samples.

Although the capillary bed can be considered homogenous at the whole cerebral length scale, heterogeneity of its spatial distribution exits. From the marmoset samples analysed by Risser *et al*. [72], the opercular visual cortex appeared strongly vascularised by capillaries (volume fraction=2.36%) in comparison with the infero-temporal cortex (volume fraction=1.60%). Geometric parameter values in this study were computed solely using the morphology of the cortical vessels in the temporal lobe due to the lack of accurate information in other regions. However, the model is not sensitive to either of the parameters (see Table 9); the impact of regional differences in the cortical capillaries on the model can be considered as minimal.

However, it has been pointed out that there is a clear distinction between the vascularisation in the grey and white matters [29]; a considerable drop in capillary density is seen in white matter with an increase toward larger vessels [73]. Such morphological difference could have a greater impact on oxygen transport. Recently, Buch *et al*. [91] introduced a method that allowed vessels with a diameter around 50 *µ*m be to imaged *in vivo*, which could help to gain further insights into the vessels in the white matter with possible quantification.

A REV was then identified based on an acceptable error margin between the parameter effective values and the voxel means. The minimum REV was found to be the 375 *µ*m cube with a 4.5% error tolerance, agreeing with the results based on perfusion simulations [12]. Results of oxygen transport simulations in discrete capillary networks also showed small differences between the 375 *µ*m and the 625 *µ*m cube [14]. Capillary cube with a side length of 375 *µ*m is therefore recommended for future studies on oxygen transport in discrete networks for it is large enough to capture the micro-scale heterogeneities yet sufficiently small compared to a typical MRI/PET voxel.

Maximum consumption rates of oxygen in the grey and white matters were optimised to achieve an OEF of 0.4. This led to an interesting oxygen distribution along the depth; an increase in oxygen level was observed at the grey-white matter interface from the 1D results. On the other hand, 1D steady state simulation from a similar model by Payne and Mai [24] showed continuous decrease in 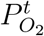 from the cortical surface to the ventricle surface. The main reason behind this difference is how oxygen is introduced to the system. While the model in this study had a constant *C*_*a*_ throughout the domain, the model in [24] applied a DBC on the cortical surface in the arteriolar compartment. Nevertheless, as discussed in Section 5.1, it is inconclusive how oxygen level changes over the depth of the brain.

Results of the 3D model showed a much wider 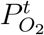 range (3–43 [mmHg]) than that of the 1D model (37–41 [mmHg]) with its lowest value found near the ventricle surface. This is mainly due to the anatomical inaccuracy of the 3D mesh leading to lower perfusion rates near the ventricles. Payne and Mai [24] employed an anatomically more accurate brain mesh and observed small variations in perfusion rates within the grey and white matters, which lead to a spacially more uniform oxygen distribution within both matters. This highlights the importance of anatomical accuracy and the potential variabilities in simulation results between individuals.

Sensitivity analysis showed that the model was not sensitive to most of its parameters. However, perturbations in oxygen solubilities and τ, the ratio resulted from linearising the ODC, had a significant impact on the tissue compartment. It is, therefore, important to accurately capture the ODC for the relevant range of blood oxygen partial pressure. Although the linear relationship used in this study could give reasonable approximations in healthy cases, it would be less suitable under pathological conditions such as ischaemia. It is also worth noting that the optimisation of the coupling coefficients [19], and subsequently the maximum consumption rates of oxygen, were performed on this patient specific brain mesh. Given the greater sensitivity of the perfusion model towards the coupling coefficient values [20], it is possible that re-optimisation may be required for different mesh geometries.

Lastly, in addition to the assumptions made while deriving the oxygen transport model, other limitations of the model itself include that the current model is purely passive. The brain is a highly regulated organ and there is a close match between blood flow and oxygen consumption. Incorporating active changes in blood flow in response to metabolic change would improve the model’s ability in analysing associated mechanisms. Furthermore, brain cells and interstitial space are considered as a whole and modelled together as the tissue compartment. Separating them into individual compartments together with the addition of mechanical responses of tissue would allow the model to capture important consequences of brain injuries, such as oedema formation after stroke. Nevertheless, the present study has demonstrated the validity of taking a porous continuum approach to model organ-scale oxygen transport; as a next step, we will examine the model’s capabilities of investigating the response of oxygen distribution to large vessel occlusions.

## Acknowledgements

SJP is supported by a Yushan Fellowship from the Ministry of Education, Taiwan (# 111V1004-2).

